# ORFLine: a bioinformatic pipeline to prioritise small open reading frames identifies candidate secreted small proteins from lymphocytes

**DOI:** 10.1101/2021.01.21.426789

**Authors:** Fengyuan Hu, Jia Lu, Manuel D. Munoz, Alexander Saveliev, Martin Turner

**Affiliations:** Laboratory of Lymphocyte Signalling and Development, The Babraham Institute, Babraham Research Campus, Cambridge, CB22 3AT, United Kingdom

**Keywords:** Lymphocytes, micropeptides, small open reading frames, Riboseq

## Abstract

The annotation of small open reading frames (smORFs) of less than 100 codons (<300 nucleotides) is challenging due to the large number of such sequences in the genome. The recent development of next generation sequence and ribosome profiling enables identification of actively translated smORFs. In this study, we developed a computational pipeline, which we have named ORFLine, that stringently identifies smORFs and classifies them according to their position within transcripts. We identified a total of 5744 unique smORFs in datasets from mouse B and T lymphocytes and systematically characterized them using ORFLine. We further searched smORFs for the presence of a signal peptide, which predicted known secreted chemokines as well as novel micropeptides. Five novel micropeptides show evidence of secretion and are therefore candidate mediators of immunoregulatory functions.

## Introduction

Open reading frames (ORFs) are the regions of the genome which contain the triplet nucleotide codons that direct the sequence of amino acids (AA) in a protein. ORFs of less than 100 codons, referred to here as small ORFs (smORFs), are particularly numerous and have been challenging to annotate and to functionally characterise (reviewed in (1–3)). smORFs have been classified according to their location relative to the main ORF within the host transcript (4). The translation products of smORFs, termed micropeptides, have been shown to be involved in many aspects of life (5–12).These new discoveries complement already characterised peptides and small proteins known to be important biological regulators. Within the immune system the best characterised of these include host defence anti-microbial peptides, chemokines and cytokines that are known to play essential roles in normal and pathological immune reactions.

The advent of next-generation sequencing technologies and proteomic approaches has led to a more comprehensive annotation of genes, transcripts and their translated protein products (3). Several large-scale genomic studies have revealed that a much larger fraction of the genome is transcribed and translated than was anticipated (13,14). A large collection of putative translatable smORFs have been identified by computational methods based on the level of DNA and protein sequence conservation across species, coding potential and context of the initiation codon. Ribosome profiling (Ribo-Seq), an approach based on deep sequencing of isolated ribosome-protected mRNA fragments, has provided extensive evidence for the translation of smORFs (14–21). A variety of metrics and algorithms can use Ribo-Seq data to annotate translated regions of the genome. Amongst them ORFScore is a metric to quantify the bias of the trinucleotide periodicity pattern of ribosome protected footprints (RPFs) towards the first reading frame in an ORF (18). The periodicity pattern has been used by several algorithms and pipelines including ORF-RATER (19), RibORF (20), RiboTaper (22), RP-BP (23), and RiboCode (24). A recently described integrated platform called RiboToolkit provides a one-step server for comprehensive analysis of Ribo-seq data and utilises RiboCode as part of its packages (25). In addition to ORFScore, other metrics can be used in conjunction to improve actively translated ORF identification. For example, the Ribosome Release Score (RRS) detects the termination of translation at the stop codon and can robustly distinguish protein-coding transcripts from ncRNAs (26).

Here we describe a new analytical pipeline that we call “ORFLine” that performs a comprehensive and systematic analysis of RNA-Seq and Ribosome profiling to identify actively translated smORFs. In comparison to previously published pipelines, ORFLine reduces computational demands by focusing on smORFs. Also, ORFline applies a series of logical filters to improve the stringency of prediction. Predicted smORFs are classified according to their host transcript type and the position of the smORF relative to other ORFs within the same transcript. We have applied ORFLine to datasets of mouse lymphocytes and discovered 5744 actively translated smORFs during B and T cell activation. We also analysed the genetic conservation, translation efficiency, and related biological processes of the predicted smORFs. We further identified smORFs containing signal peptides which have a potential to be secreted and could act as immune regulators.

## Results

### Overview of ORFLine

ORFLine takes Ribo-Seq, RNA-Seq, reference genome, transcriptome and gene annotation as input data and produces an output list of predicted smORFs with genomic coordinates and classification (**Figure 1A**). The three main pipeline components to process the raw Illumina sequences and perform smORF prediction are: 1) Prediction of putative smORFs; 2) Sequencing data QC and processing; and 3) Identification of translated smORFs. Prediction of putative smORFs and sequencing data processing are independent components and can be executed in parallel. The identification of translated smORFs utilises the output of the previous two components as input (**Table 1**). ORFLine is applicable to data from any species.

**Fig. 1.**
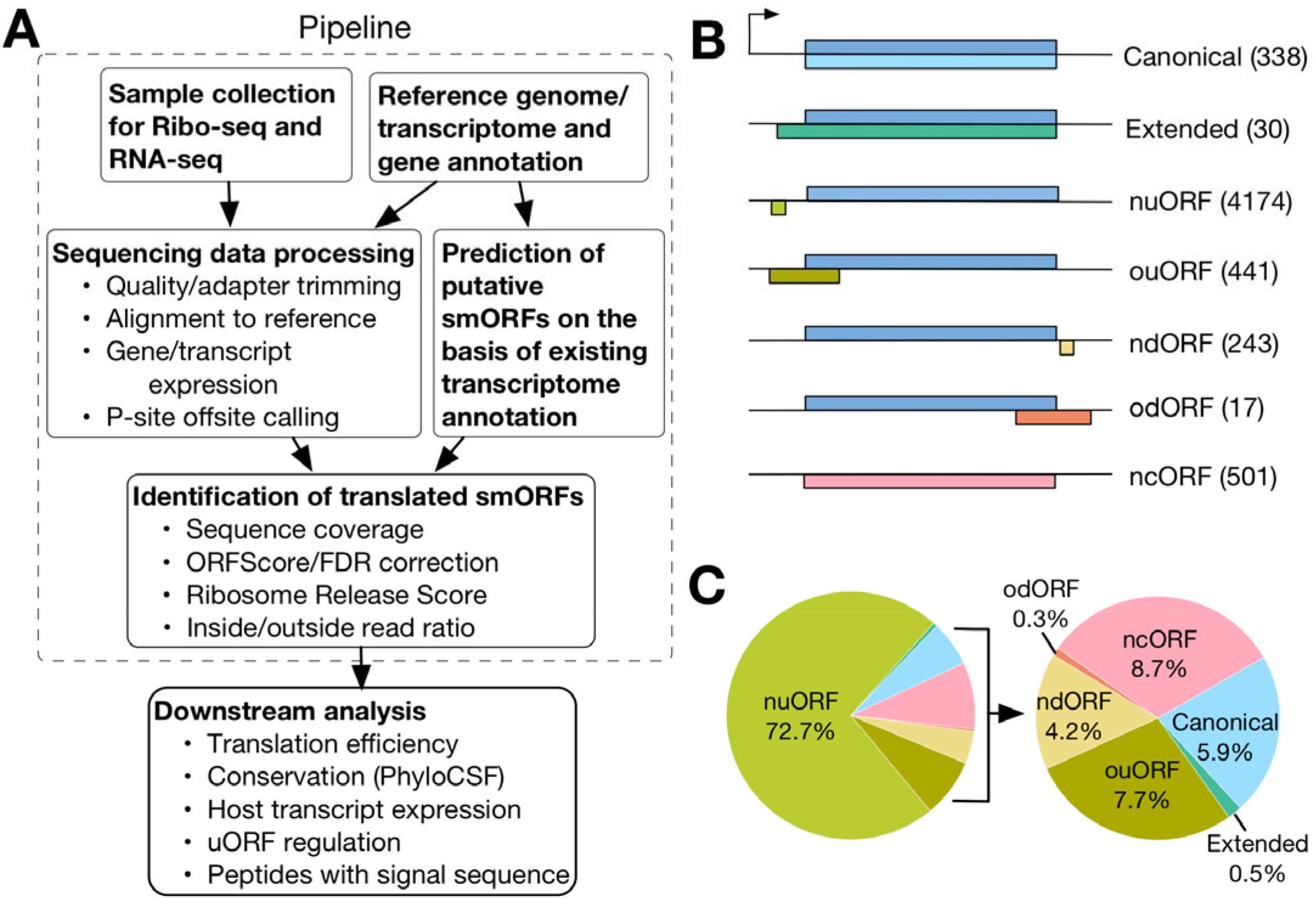
Identification of different classes of actively translated smORFs in this study. **(A)** Computational pipeline (in dashed line square) to identify translated smORFs. Sequencing data for RNA-Seq and Ribosome profiling is processed and the reads mapped to the mouse reference genome GRCm38/mm10. In parallel, putative smORFs were predicted by scanning the mouse transcriptome. Several experimental metrics for each putative smORF were quantified and the smORFs exceeding a threshold for each metric were kept for downstream analysis. **(B)** Predicted smORFs were classified into 7 groups according to their relative location in the host transcript. The number of smORFs in each class is shown in parentheses. **(C)** Pie chart showing the proportion of smORFs of different classes.

**Table 1.**
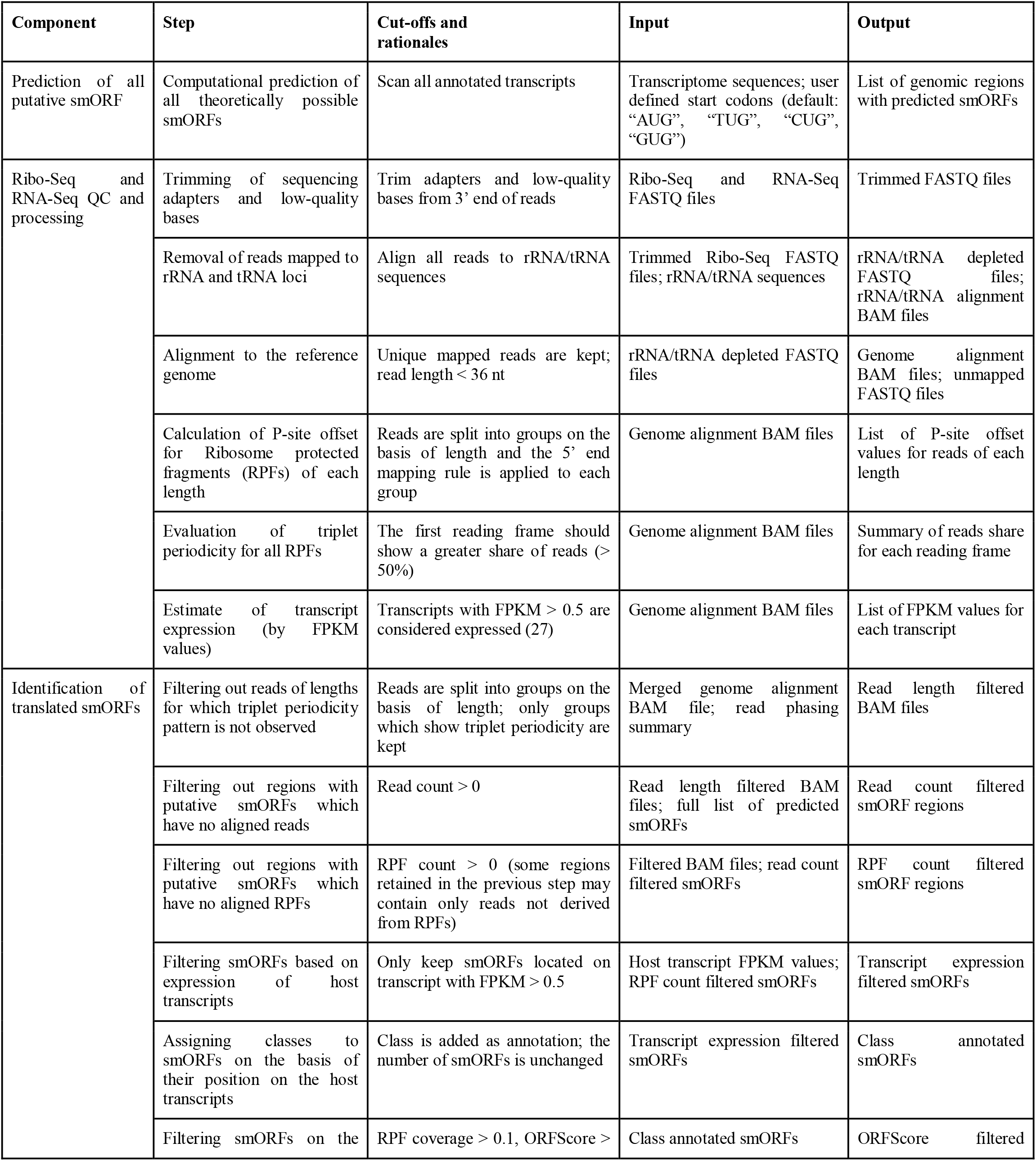
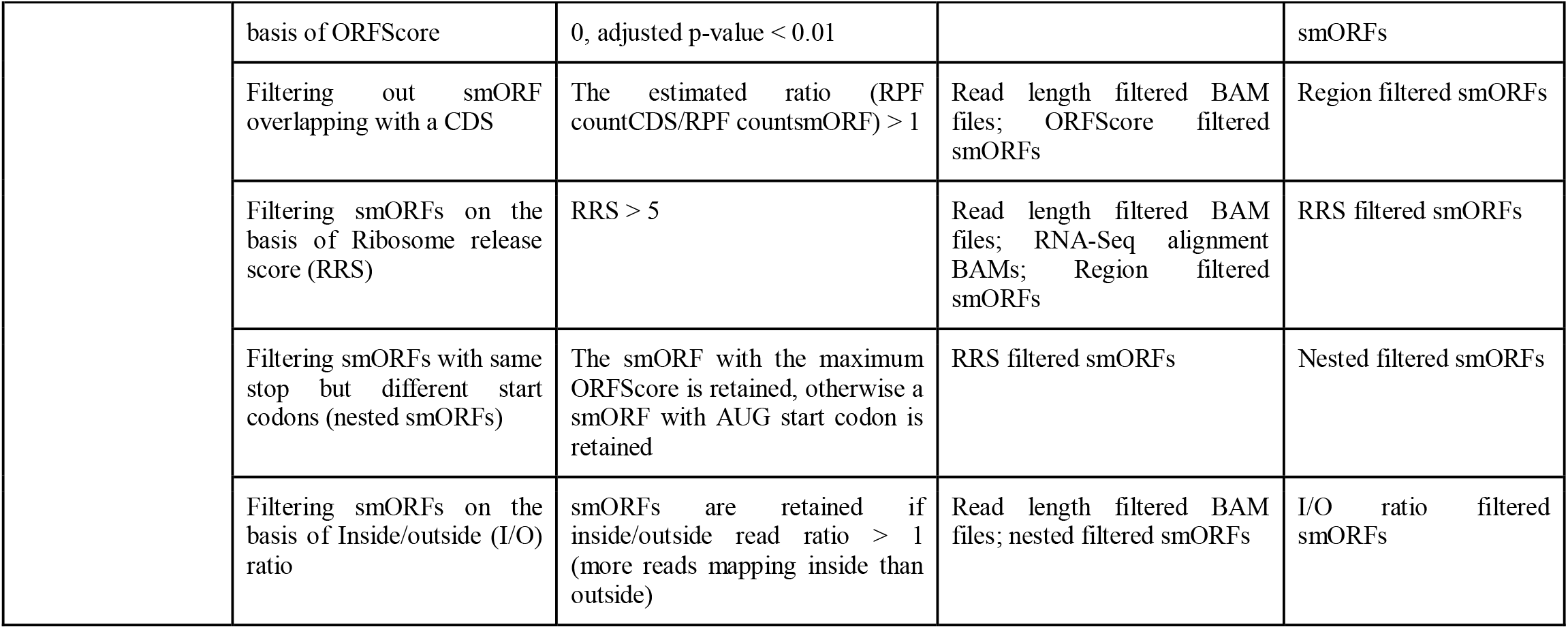
Summary of steps in the ORFLine.

The output of ORFLine is a list of smORFs that have passed the filters in the identification of translated smORFs. They are identified as smORFs with ribosome protected RNA fragments (RPF) signal. The output file (**Table 2**) contains the genomic location and splicing information (including number of exons and exon lengths) of a smORF are clearly annotated and can be loaded and visualized in a genome browser. The quantitative information about a smORF is also calculated including translation efficiency, RNA expression and Ribosome- protected RNA expression (FPKM value). The nucleotide sequences are retrieved and translated into amino acid sequences and presented in column 25 of **Table 2**.

**Table 2.**
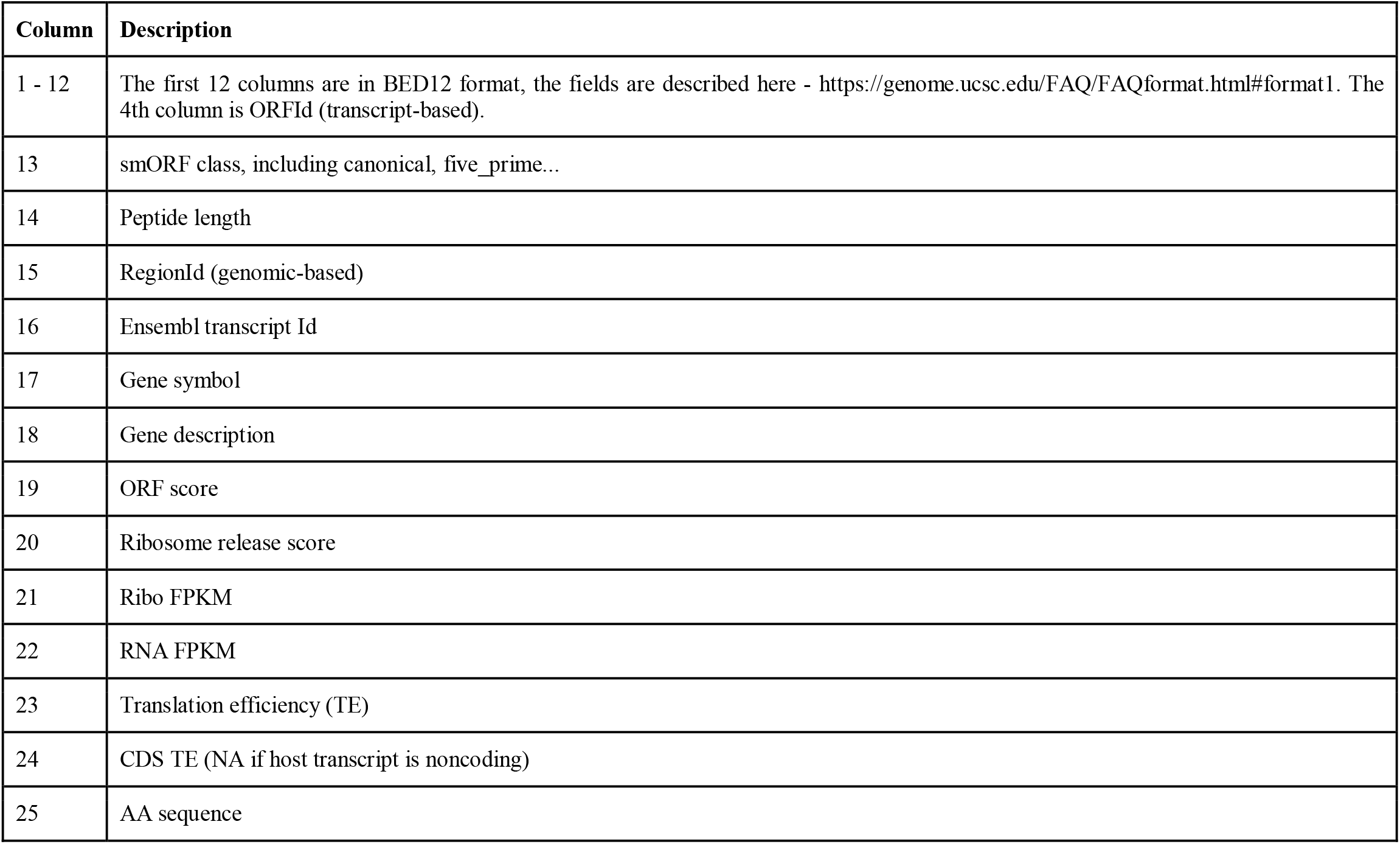
Pipeline final output format.

### Comparison between ORFLine and RiboCode

We characterised smORFs in mouse lymphocytes using datasets obtained from *ex vivo* non-activated B cells; a published dataset from our lab of Lipopolysaccharide (LPS)-activated B cells (28); a new dataset of B cells activated with LPS plus interleukins IL-4 and IL-5 for 48 hours, by which time most cells had divided once, and some started a second division; naïve CD4+ T cells stimulated with antibodies to CD3 and CD28 which mimics activation by antigen; and a published time-course of Th1 T cells re-stimulated with anti-CD3+anti-CD28 (29) see **Table S1**. ORFLine predicted a total of 5744 unique smORFs in all samples analysed (union of 2607 smORFs predicted in B cells and 4935 smORFs predicted in T cells) (**Table S3**). A lower number (568) of smORFs were predicted for the resting B cells than for LPS-activated B cells (2444), most likely reflecting an overall increase in RNA abundance associated with elevated rates of transcription in activated B cells.

We also analysed the same datasets with the recently published ORF- detection pipeline RiboCode (24) using its default settings. RiboCode predicted a total of 15,920 unique smORFs, of which 3,667 are smORFs nested in longer smORFs in the same reading frame and 48 smORFs are from non-expressed transcripts. We removed those 3,715. In the remaining 12,205 smORFs, 3,337 were predicted as internal or frameshift smORFs. These are found nested in the CDS, but in a different reading frame. Considering that frameshift translation is a rare event (30), they are not included in our results. We removed all 3,337 frameshift smORFs predicted by RiboCode and compared the remaining 8,868 non-internal smORFs predicted by RiboCode with the 5,744 predicted from ORFLine (**Figure S2**). Of these, 1,957 (22.1% in RiboCode and 34.1% in ORFLine) are found as exact genomic coordinate matches by both pipelines (**Table 3**). For un-annotated smORFs, we are not certain they are translated, and we lack a reference set of true-positives, therefore we sampled the smORFs which are differentially identified by the two pipelines and noticed that smORFs predicted by RiboCode typically have low RPF coverage or are assigned a low or negative ORFScore, or low RRS, and are filtered out by ORFLine (**Figure S3**). Our criteria for metrics have shown to be robust in smORF prediction in previous studies (18,26). ORFLine also predicted 356 smORFs encoded by low abundance transcripts (25% percentile) that are not predicted by RiboCode. Taken together, these comparisons demonstrate that ORFline allows a reliable identification of smORFs residing in transcripts of low abundance.

**Table 3.**
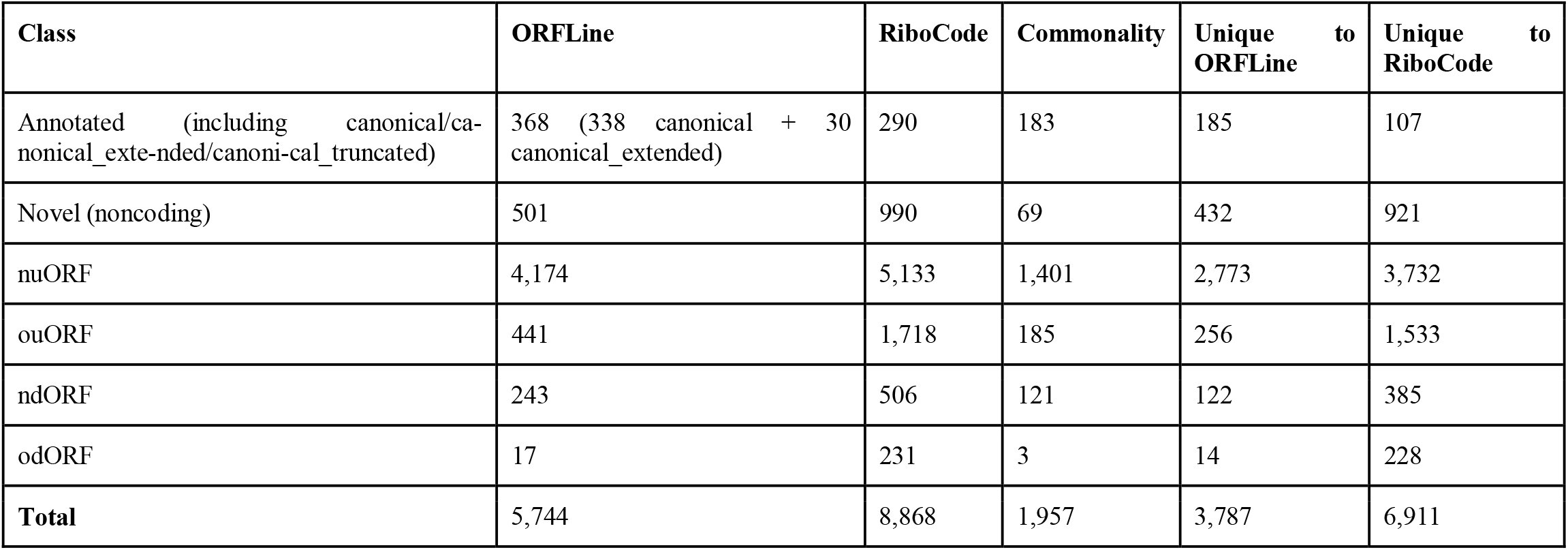
ORFLine and RiboCode prediction commonality/difference by class.

### smORF classification

ORFLine classified smORFs according to their relative position with the annotation, if any, of the host transcript (**Figure 1B**). It predicted canonical smORFs and extended variants of annotated coding DNA sequences (CDS) of 100 codons or less in protein-coding mRNAs. ORFLine also found upstream ORFs (uORFs) which we subdivided into either uORFs starting in the 5’ untranslated region (5’ UTR) of annotated protein-coding mRNAs and overlapping with the coding region (ouORFs) or non-overlapping uORFs (nuORFs) which terminated before the start of the annotated CDS. uORFs are known to be prevalent in the genome and they represent 80% of all smORFs predicted by ORFLine (**Figure 1C**). In addition, ORFLine identified downstream ORFs (dORFs) as the rarest class of smORFs. These can be subdivided as overlapping dORFs (odORFs) that overlap with the CDS and extend into the 3’UTR of known protein-coding mRNAs or non-overlapping dORFs (ndORFs) located exclusively in annotated 3’UTRs. Lastly, 501 smORFs in putative non-coding RNAs (long noncoding RNAs and pseudogenes) were predicted, which are termed ncORF. Direct biochemical and functional evidence is available for only ~40% (338) of canonical smORFs in protein databases such as UniProt (The UniProt Consortium, 2019) for their protein products, it includes diverse entities such as chemokines and subunits of mitochondrial complexes. The remainder (~5340) have either not been functionally characterised or have not been annotated at all.

### smORF conservation

To examine the conservation of smORF-encoded micropeptides between species, we employed PhyloCSF to analyse signatures of evolutionary conservation. 11.4% of smORFs had a PhyloCSF score > 50, thus showing strong evidence of conservation (**Figure 2A**), with canonical smORFs being enriched among these (**Figure 2B**). A small subset (~6.5%) of uORFs, ncORFs and dORFs showed high PhyloCSF scores, indicating conservation of smORFs CDS. There are over 60% of smORFs lacking signs of selective pressure to maintain their amino acid sequences (no cross-species sequence alignment and not conserved, **Figure 2A**), in which uORFs, ncORFs and dORFs are enriched (**Figure 2B**). The median length of canonical smORFs is 79 codons, however, the median length of uORF, dORF and ncORF are 24, 34 and 33 codons respectively. By comparison with other classes, canonical smORFs are, on average, longer and more highly conserved (**Figure 2C**). Having distinct transcript organization, size, conservation and peptide structure, the cellular and molecular functions of canonical smORFs, uORFs, dORFs and ncORFs are likely to differ from each other, with less conserved classes primarily independent of peptide sequences. However, we observed that the PhyloCSF score positively correlates with the length of coding sequence (data not shown). Therefore, it is likely that the conservation of shorter smORFs is underestimated.

**Fig. 2.**
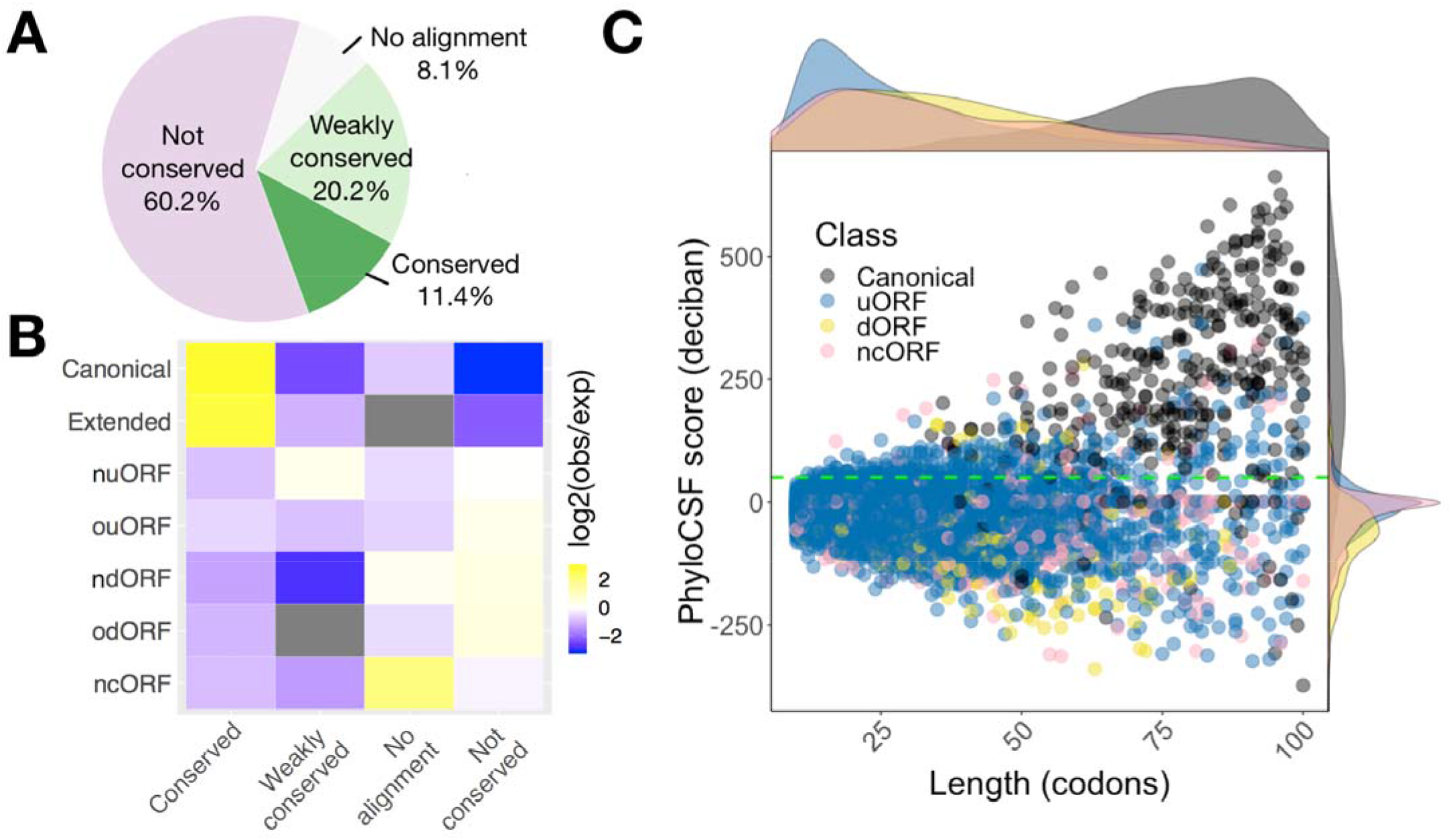
smORFs showing different conservation and length distributions according to their classes. **(A)** Most smORFs are not conserved at the peptide level. Pie chart represents the coding potential (PhyloCSF score). smORFs with PhyloCSF score ≥ 50 are considered conserved. smORFs are considered weakly conserved if their PhyloCSF scores are positive but smaller than the threshold 50. **(B)** Canonical and extended smORFs are enriched in conserved peptides. Enrichment heatmap depicts log 2 ratio of the number of smORF observed (obs) to the number of smORF that would be expected (exp) by chance given overall distributions of smORF classes and conservation levels. **(C)** Scatter plot shows the distributions of codon length and PhyloCSF score for each smORF type. Marginal densities of length and PhyloCSF score are also shown on the top and the right-hand side of the scatter plot. Green dashed line indicates a PhyloCSF score of 50. Here the original classification in Fig 1B was simplified by combining the canonical and canonical extended ORFs as canonical; nuORF and ouORF as uORF; and ndORF and odORF as dORF. Canonical smORFs are on average longer and more conserved than other types.

### Canonical smORFs

A total of 338 canonical smORFs were predicted in B and T cells. 88% of these are conserved or weakly conserved between species (**Figure 3A**). We divided canonical smORFs into “short CDS” and “short isoforms”, the latter are the products of alternative splicing of transcripts from genes annotated as encoding proteins longer than 100 amino acids (4). Among the predicted canonical smORFs, 54.4% are short CDSs and 45.6% are short isoforms. There are hundreds of putative short CDSs in mouse and human, these are typically located on monocistronic transcripts and their host transcripts are structurally shorter and simpler compared with canonical mRNAs (4). We have predicted 184 short CDSs and they have a median size of 79 codons. We find short isoforms have a median size of 80 codons and resemble short CDSs in size and conservation (**Figure 3B**). As short isoforms share conserved protein sequences with their longer canonical protein isoforms, they may have functions that are directly related to their longer protein isoforms (4).

To increase confidence that predicted canonical smORFs were indeed translated we calculated the translation efficiency of the short CDSs and short isoforms. When compared to long CDSs of expressed proteincoding transcripts, we found their median translation efficiency to be greater (**Figure 3C**). We also conducted GO term enrichment analysis comparing the 184 short CDS and the 154 short isoforms against the remaining 3536 smORF-encoding genes of B and T cells. The top hits of short CDS are related to chemokine activity and mitochondrial biology (**Figure 3D, Table S4**). Seven chemokines are predicted (Ccl1, Ccl22, Ccl3, Ccl4, Ccl5, Cxcl10, Cxcl11). We also observed enrichment of gene products involved in mitochondrial complexes, for example, Uqcr10 is a subunit of Coenzyme Q: cytochrome c reductase (Complex III); this complex has a critical role in the oxidative phosphorylation pathway for the generation of ATP. Another mitochondrial protein is Romo1, which is located in the mitochondrial membrane and is responsible for increasing the level of reactive oxygen species (ROS) in cells (31). Romo1 also has antimicrobial activity against a variety of bacteria by penetrating the bacterial membrane (32). Short isoform encoding genes are associated with a broad range of GO terms, with no single term for GO biological processes reported as enriched.

### uORFs

Approximately 50% of annotated animal mRNAs contain uORFs (4,33) and translation of uORFs has been widely reported in different organisms (21,34,35). We observed that the median translation efficiency of uORFs is greater than that of long CDS (**Figure 4A**). About 4% of the uORFs have a high PhyloCSF score and TE (above the median TE of long CDS) and potentially encode conserved functional micropeptides (**Figure 4B**, for B cells activated with LPS and IL-4+IL- 5). However, the sequences of the majority of uORFs are not conserved, suggesting that any potential function is largely independent of the encoded peptide. The proportion of expressed uORF-containing transcripts in B cells and T cells is between 6.2% and 12.4%, except resting B cells (2.7%). It has been demonstrated that uORFs may regulate the translation of the downstream CDS. Several studies have shown a repressive effect of uORFs on the translation of CDS (21,36,37). We analysed the effect of uORFs on mRNA translation of their associated genes by comparing the translation efficiency of the CDS in all uORF-containing transcripts versus those lacking uORFs. As expected, the presence of uORFs and overlapping uORFs was associated with a translation repression (**Figure 4C**). We performed GO enrichment analysis for all uORF-containing genes to discover their associated biological processes (2881 target genes against 3536 background genes) and these genes are mostly enriched in processes linked to protein modification, regulation of gene expression and cellular response to stimulus (**Figure 4D, Table S4**). This indicates that uORF-containing genes are broadly involved in complex biological pathways such as protein or RNA production and cell signalling. Regulatory uORFs may be suited to allow the rapid changes in gene expression in response to stress and environmental stimuli.

**Fig. 4.**
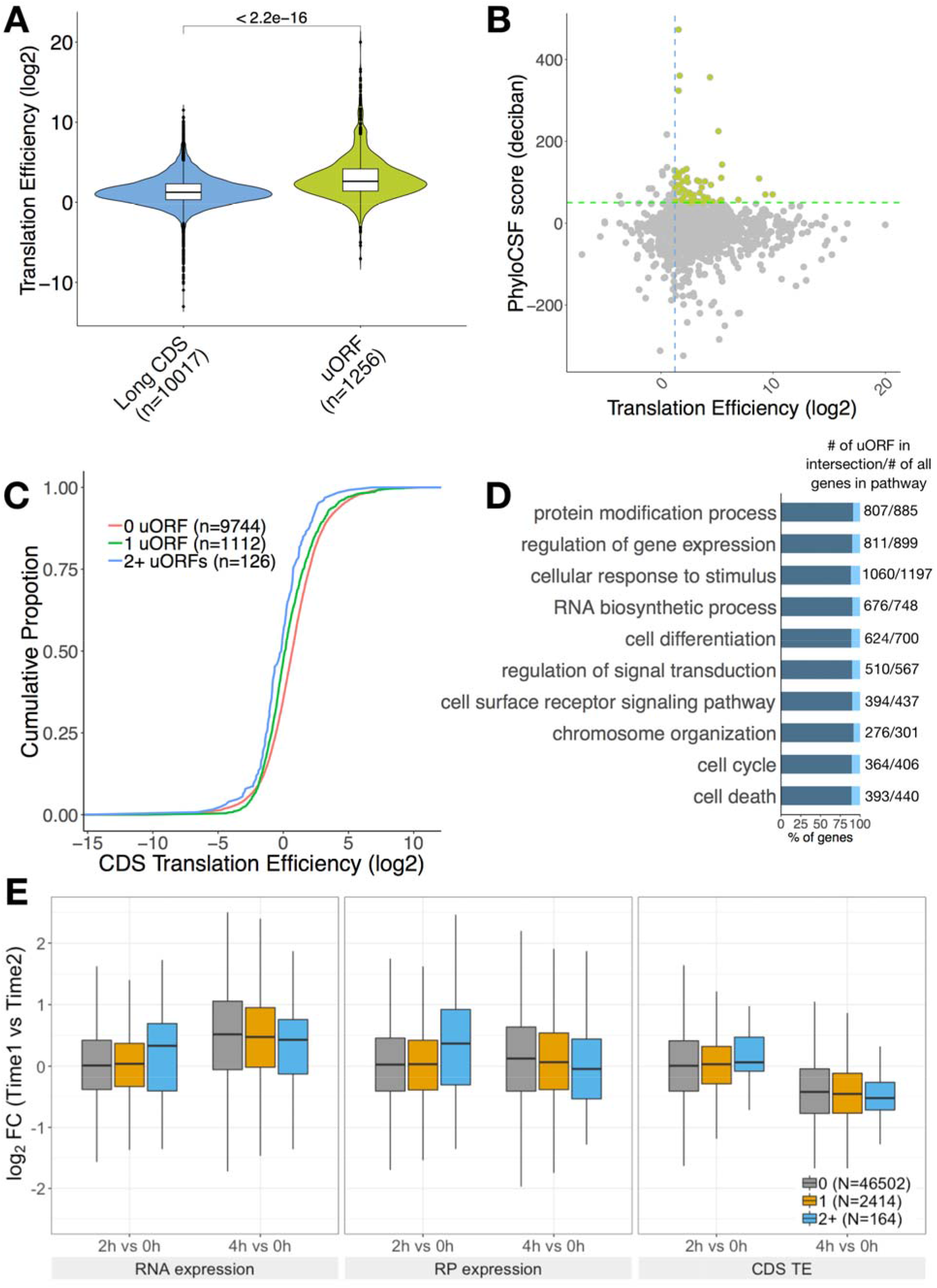
uORFs regulate the translation of their downstream CDS. **(A)** Translation efficiency distributions of long CDS and uORF. Significance was computed using a two-sided Mann-Whitney test. **(B)** Scatter plot of uORF translation efficiency and PhyloCSF score. Green broken line represents a PhyloCSF score of 50 used as a threshold for conservation, blue broken line represents the median TE of long CDS. uORFs that are conserved and have a high TE are highlighted. **(C)** Cumulative distribution of translation efficiency in expressed uORF-containing transcripts versus transcripts lacking uORFs as control. Significance was computed using two-sample Kolmogorov–Smirnov test for each uORF set compared to the control. (1 uORF P = 1.321e-14, 2+ uORFs P = 1.828e-6). **(D)** Biological process gene ontology terms found to be significantly enriched in the uORF-containing gene list. **(E)** Translational regulation of downstream CDSs by uORFs during T cell activation. Logarithmically transformed fold change of CDS RNA abundance (FPKM), RPF abundance (FPKM) and TE between two time points (2h vs 0h and 4h vs 0h) in 0,1, 2+ uORF-containing transcripts. Significance was computed using a two-sided Mann-Whitney test. For RNA expression, there is no statistically significant difference between 0 and 1, 0 and 2+ or 1 and 2+ uORF-containing transcripts for 2h vs 0h (P= 0.4717, 0.4399 and 0.491, respectively) and 4h vs 0h (P= 0.3169, 0.0542 and 0.1057, respectively). For abundance of RPFs, the comparisons between 0 and 1, 0 and 2+ or 1 and 2+ uORF-containing transcripts are: 2h vs 0h, P= 0.7073, 0.2558 and 0.2658, respectively; 4h vs 0h, P= 0.05284, 0.04256 and 0.1409, respectively. For CDS TE, no statistical significance was observed when comparing 0 and 1, 0 and 2+ or 1 and 2+ uORF-containing transcripts in 2h vs 0h (P=0.5685, 0.3602 and 0.425, respectively) or 4h vs 0h (P=0.2296, 0.2481 and 0.4238, respectively). Outliers are not displayed.

We then investigated the influence of uORFs on translation of the downstream CDS during the first four hours of T cell activation **(Figure 4E)**. During the first two hours of activation the median RNA and RPF abundance for both non-uORF-containing transcripts and for transcripts containing a single uORF remains mostly unchanged as the median Log2 fold change is close to 0. There were few transcripts with two or more uORFs and these had a median 0.35 Log2 fold change of RNA and RPF abundance that is not statistically significantly different from the other two groups. However, we cannot rule out that we would find a difference if the numbers of transcripts in this class was greater.

By four hours all mRNAs, irrespective of the presence of uORFs, show a median Log2 fold change of RNA abundance of around 0.5 compared to 0 hours. At 4h RPFs did not increase substantially, indicating a possible delay of translation of induced transcripts. As transcripts with two or more uORFs show a negative and the lowest median Log2 fold change of RPF abundance at 4h compared to 0h (P=0.04256 when compared to non-uORF-containing transcripts) the presence of uORFs appears to have a negative impact on translation of the downstream CDS. For translation efficiency, there is little change during the first two hours. The negative median Log2 fold change in TE for all groups in 4h vs 0h reflects the increase of RNA abundance at 4h and the lack of change of transcripts in RPFs. These results indicate that the rate of translation lags behind the increase in transcript abundance and the presence of uORFs can affect translation in activated T cells.

### smORFs in non-coding RNAs

Non-coding ORFs (ncORFs) are smORFs that are found in annotated long non-coding RNAs (lncRNAs) and pseudogenes. They are typically short with a median length of 33 codons. By definition, non-coding RNAs are not translated into protein. However, annotated lncRNAs have been predicted from their sequences to contain six smORFs on average (4). We have predicted 501 translated ncORFs and 14.4% of these are considered conserved or weakly conserved. We noticed very different distributions of size and PhyloCSF score between ncORFs and canonical smORFs (**Figure 5A**). The distribution of translation efficiency for ncORFs is also different from that for long CDS, the median TE of ncORFs is greater than long CDS (**Figure 5B**). Three ncORFs identified in LPS-activated B cells (Cct6a, Gm16675 and 6330418K02Rik) were found to have a high PhyloCSF score and TE, so we infer them to encode functional micropeptides (**Figure 5C**). We searched the micropeptides they encode in NCBI BLASTp database (38), but did not find any match for Gm16675. The 6330418K02Rik gene is annotated as an antisense lncRNA gene in GENCODE and only one match was reported for its predicted micropeptide (sequence ID: EDL19200.1). The micropeptide was partially aligned to three uncharacterized proteins with 35.59% to 78.18% identity in *Habropoda laboriosa* and *Gulo gulo.* The examples likely reflect that these genes are misclassified as non-coding, although it is possible that they could also have functions as a noncoding RNA in addition to their peptide coding capacity.

**Fig. 5.**
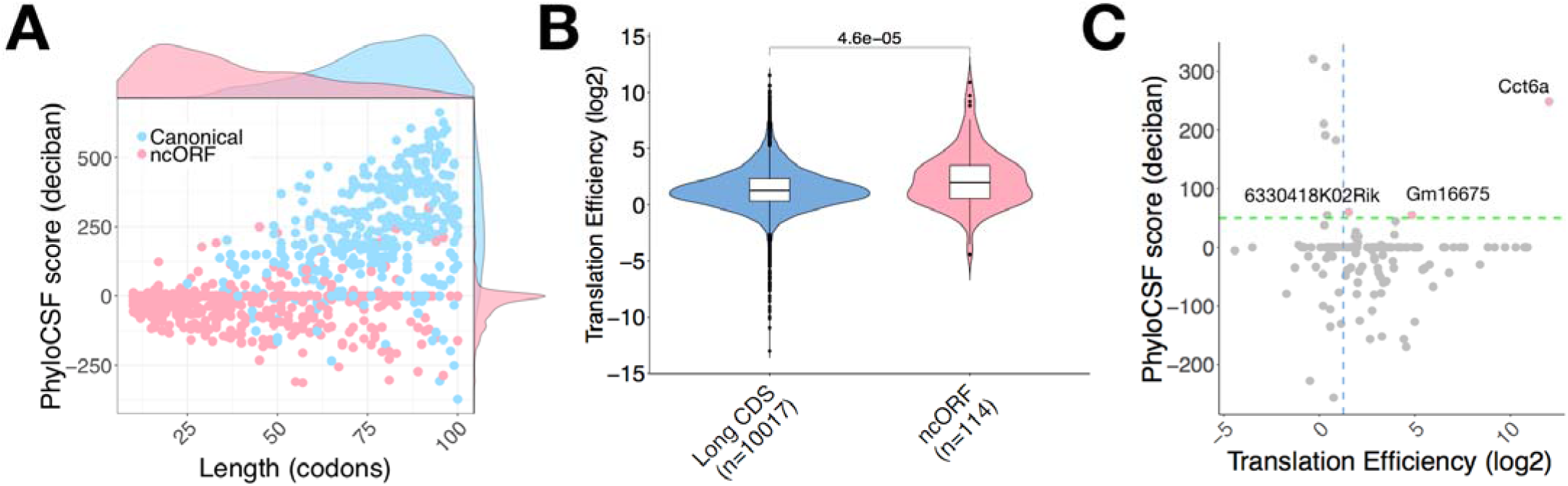
Translated smORFs predicted in noncoding RNAs. **(A)** Canonical smORFs and ncORFs showing very different distributions in length and PhyloCSF score. **(B)** Translation efficiency distributions of long CDS and ncORF. Significance was computed using a two-sided Mann-Whitney test. **(C)** Translation efficiency and PhyloCSF score are shown for ncORFs (LPS-activated B cells). Scatter plot of ncORF translation efficiency and PhyloCSF score. Green broken line represents a PhyloCSF score of 50 used as a threshold for conservation, blue broken line represents the median TE of long CDSs. ncORFs that are conserved and have high TE are highlighted. Three genes (Cct6, 6330418K02Rik, Gm16675) potentially encode micropeptides.

### dORFs

243 ndORFs and 17 odORFs were predicted with a median length of 34 AA. Only 20 (~7.7%) are conserved or weakly conserved (**Table S5**). The translation efficiency of dORFs is lower than the long CDSs in general (data not shown). In transcripts that contain multiple ORFs, a translation reinitiation mechanism is able to prevent recycling of some or all ribosome subunits upon termination of the first translated ORF and thereby enable the translation of the dORF (39). The low TE indicates a very low level of translational reinitiation after the stop codon of the upstream CDS. In B cells activated with LPS and IL-4+IL- 5 we noticed that dORF-containing transcripts show no significant difference in TE compared to those without. (**Figure 6**).

**Fig. 6.**
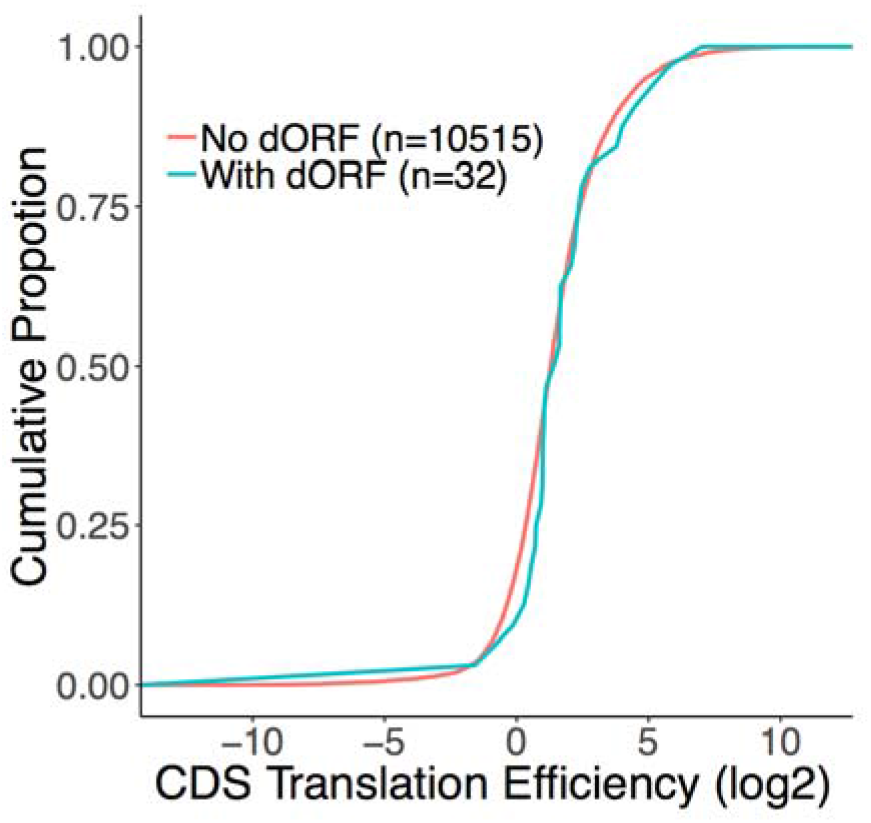
The canonical ORFs of dORF-containing transcripts may be translationally enhanced. Cumulative distribution of translation efficiency in expressed dORF-containing transcripts versus transcripts lacking dORFs as control. Significance was computed using a two- sample Kolmogorov–Smirnov test, P = 0.437.

### Signal sequence containing micropeptides

An N-terminal signal peptide sequence of 16-30 amino acids is characteristic of proteins destined to be secreted, resident within cellular membranes or within compartments of the secretory pathway. We predicted the presence of signal peptides in amino acid sequences of micropeptides using SignalP server (40). This predicted 80 candidates including known chemokines (CCL-1, −2, −4, −5 and −22) and the cell surface protein CD52, as well as the recently identified lncRNA encoded Aw112010 (12) and 1810058I24Rik micropeptides (41). Amongst these, 28 are canonical micropeptides and they typically have high levels of conservation. Of the remaining 52 non-canonical micropeptides, 12 show conservation (**Figure 7A**).

**Fig. 7.**
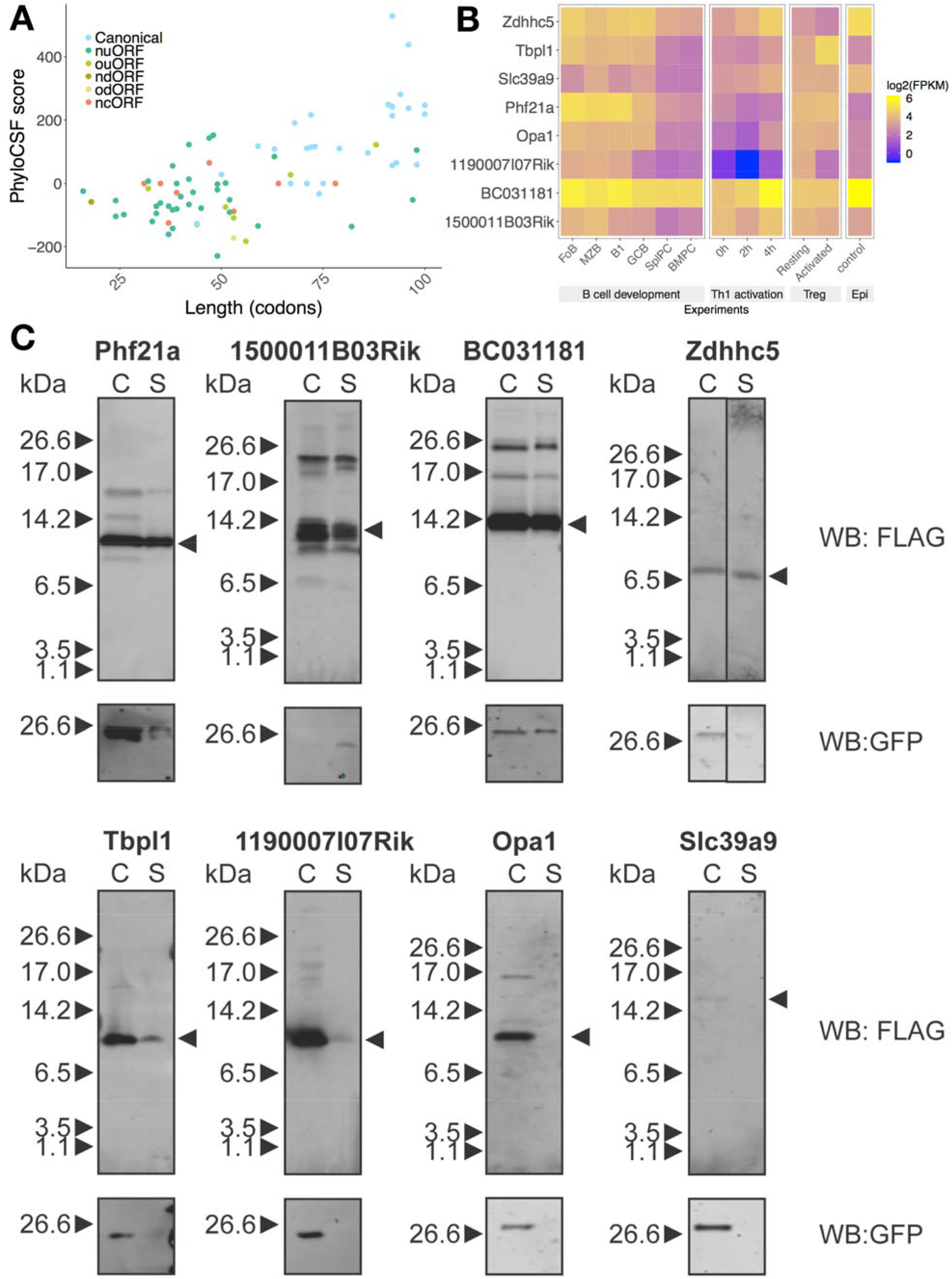
Predicted signal sequence containing micropeptides and their host transcripts expression under different conditions. **(A)** Scatter plots show the distributions of length (codon) and PhyloCSF score for each predicted signal peptide containing micropeptides. **(B)** Heatmap analysis of host transcript expression during B cell terminal differentiation, Th1 cell activation, resting/activated regulatory T cells and epidermal cells (Epi). Selected micropeptides are shown in the heatmap, they are conserved in humans and there is limited or no information regarding their function. They are ordered by length. **(C)** Expression and secretion of micropeptides. Plasmids encoding predicted micropeptides were transfected into 293T cells and micropeptides in total cell lysate (C) and total secreted fraction (S) were detected by anti-FLAG antibody. GFP expression indicates transfection efficiency.

We selected eight putative smORFs for further characterisation (**Figure 7B**). First of all, except for the uORF from Zdhhc5, all of uORF-encoded micropeptides are in different reading frames from the main CDS. The coding sequence of Zdhhc5 uORF does not overlap with the ZDHHC5 ORF, suggesting it is not an N-terminal extension of the main CDS. To examine the expression of signal sequence-containing micropeptide host transcripts, we compared mouse RNA-Seq datasets for lymphocytes spanning B cell terminal differentiation (42), Th1 cell activation (29), and regulatory T cells (43) as well as epidermal cells (44). These data revealed dynamic expression patterns for several of the host transcripts. For example, BC031181 was downregulated during B cell differentiation but upregulated during Th1 cell activation, it was also highly expressed in epidermal cells (**Figure 7B**). Host transcript expression patterns provide a lead to where and what stage of cell differentiation micropeptides may be produced and can help with experimental validation of micropeptide prediction. To determine if the selected putative micropeptides are likely functional, we compared the conservation of amino acid sequence between different mammalian species (**Figure S4**). All of the micropeptides including those encoded in uORFs show evidence of conservation. This indicates positive selection pressure of the coding sequence of these micropeptides.

### In vitro characterisation of predicted micropeptides with signal sequence

We sought to validate the potential for secretion of micropeptides with signal sequences. To this end, we selected and cloned eight putative smORFs with predicted signal sequences into a dicistronic mammalian expression vector which allowed synthesis of the micropeptide with a FLAG epitope tag at its C-terminus and of GFP driven by an IRES from the same transcript. HEK293T cells transfected with micropeptide-encoding plasmids displayed anti-FLAG signals in both total cell lysates (C) and supernatant (S) fractions (**Figure 7C**). GFP was detected in all total cell lysates, but the smORF encoded in the Slc39a9 transcript showed no evidence of expression. The smORFs encoded by Phf21a (uORF), 1500011B03Rik and BC031181 (both canonical smORFs) showed the most abundant expression and secretion. The small ORFs encoded by Zdhhc5 and Tbpl1 expressed less strongly but showed evidence of secretion. By contrast, the smORFs encoded by Opa1 and 1190007I07Rik were weakly or not at all secreted (**Figure 7C**). A recent report demonstrates that the human ortholog of 1190007I07Rik, named *C12orf73*, encodes a functional micropeptide named BRAWNIN found at the inner mitochondrial membrane and required for respiratory complex III assembly (45). These results demonstrate that putative smORFs can be expressed and secreted, but further investigations are required to validate their subcellular localisation and to demonstrate their biological functions.

## Discussion

In this study we have developed an improved pipeline for discovery of novel smORFs expressed with low abundance. We used this pipeline to predict 5744 unique smORFs that show evidence of being translated in B and T lymphocytes in different conditions. Apart from 368 annotated as short CDSs or isoforms, the others are novel and located in long noncoding RNAs, pseudogenes or in the 5’UTR and 3’UTRs of canonical protein coding transcripts. By assessing the conservation of the amino acid sequences and their translation efficiency relative to long proteins we inferred whether the translation products of these smORFs were likely to be functional.

We compared our pipeline with RiboCode which assesses the triplet periodicity of RPFs in an ORF with modified Wilcoxon signed-rank test and is claimed to outperform other existing pipelines including RiboTaper, Rp-Bp and ORF-RATER (19,22,23). Ribocode has been incorporated into a recently published integrated tool (RiboToolkit) to analyse ribosome profiling data (25). When compared with ORFLine, RiboCode predicted more putative smORFs, some of which appeared to be ORFs with the same stop codon but different start codons. RiboCode maps Ribo-Seq reads to the transcriptome and can lead to reads being mapped to multiple transcripts potentially increasing the number of positive signals. By contrast, ORFLine maps reads uniquely to the genome. We also observed that for the different putative smORFs predicted by RiboCode and ORFLine, those unique to ORFLine have higher RPF coverage and ORFScore. Thus, ORFLine produced fewer outputs and additionally provided more diverse classification of ORFs according to positional information.

Among the predicted smORFs, 80% were located within 5’UTRs. Many of these were not conserved at the amino acid level between species and may be regulatory. A recent study has shown that non-canonical Hoogsteen-paired G-quadruplex (rG4) structures present upstream of some uORFs promote 80S ribosome formation on start codons, causing inhibition of translation of the downstream CDS (46). Identification of rG4 motifs in an uORF context thus may help to distinguish regulatory uORFs and this feature could be incorporated into new iterations of ORF calling pipelines.

About four percent of our predicted smORFs are dORFs, which is consistent with previously reported data (20). dORFs of less than 100 aa are poorly characterised. Our pipeline predicted three dORFs encoded by Pgs1, Szrd1 and Dpm2 genes which are conserved by amino acid sequences between human and mouse (data not shown). These dORFs likely encode functional proteins and deserve further investigation. Recently, Wu et. al. also reported that dORFs are poorly conserved in human and zebrafish genomes, but are readily translated and can enhance translation of the main CDS (47). Our results also demonstrate no evidence of a statistically significant effect of dORFs on translation of the upstream CDS. However, it is important to point out that we considered only 32 dORFs in LPS-activated B cells, compared with 1406 for human and 1153 for zebrafish by Wu et. al. and therefore our analysis of TE may be underpowered to detect an effect.

We predicted 8 micropeptides with signal peptide sequences and five were found to be secreted in a model system. Little is known regarding the abundance or stability of these micropeptides in physiological settings. In our attempts to validate the predictions of novel smORFs, we observed that overexpression of smORFs in mammalian cells yielded varied levels of expression. Specifically, the uORF of Slc39a9 did not produce any detectable proteins despite codon optimisation, indicating possible short half-life of this smORF encoded protein. It has been suggested that many peptide products are selectively and rapidly degraded within cells, and hence are difficult to detect biochemically (48,49). These factors impede their identification by regular mass spectrometry as they are often lost during sample preparation thus not available for detection. Recently it was proposed that immunopeptidomics based on the repertoire of peptides presented by MHC class I molecules may be suitable for detection of smORFs translation products (50). The immunopeptidome differs from the proteome in that it skews away from abundant gene products, enriching peptides from non-canonical translation initiation and micropeptides with short half-lives (51).

As the ability to predict smORFs far outstrips the ability to validate them experimentally only a small number of predicted smORFs have so far been validated. For further investigation of the biological functions of the potentially secreted micropeptides, investment into the generation of antibodies and model organisms will be required. For micropeptides with signal sequences, they have the potential to be novel cytokines. If so, it will be exciting to validate the existence of receptors and to shed light onto the biology of these micropeptides.

## Materials and Methods

### Tissue culture

B lymphocytes from the spleen or lymph nodes (LNs) of C57BL/6 mice were isolated using the B Cell Isolation Kit (Miltenyi Biotec). For activation, B cells were cultured for 48 hours in RPMI 1640 Medium (Dutch Modification) supplemented with 10% FCS, 100 IU/ml penicillin, 100 μg/ml streptomycin, 2 mM L-GlutaMAX (Gibco), 1 mM Sodium Pyruvate and 50 μM ß-mercaptoethanol in the presence of 10 μg/ml of LPS (Sigma, E. Coli 0127: B8), 10ng/ml of IL4 and 10ng/ml of IL5. Naïve CD4+ T lymphocytes from spleen and LNs were isolated with CD4+CD62L+ T Cell Isolation Kit (Miltenyi Biotec) and stimulated in the same medium as for B cells using plate bound anti-CD3 (2C11) and 1 μg/ml of anti-CD28 (37.51) for 24 hours. HEK293T cells were maintained in DMEM (Gibco) with 10% FBS (Gibco) and 1 × GlutaMAX-I (Gibco).

#### cDNA Library preparation

RNA-Seq libraries were generated using the TruSeq Stranded mRNA Sample Prep_TM_ Kit (Illumina Inc). Ribo-Seq libraries were prepared using the ARTseq™ Ribosome Profiling Kit (Epicentre, Illumina) from cells treated with cycloheximide (CHX, 100 μg/ml) prior to cell lysis. cDNA libraries were sequenced using Illumina HiSeq1000, Illumina HiSeq2000 or Illumina HiSeq2500 system in a 100-bp single-end (RNA-Seq) or 50- bp single-end (Ribo-Seq) mode. Summary metrics of libraries used are described in **Table S1**.

### Reference genome, transcriptome and annotation

GENCODE (52) reference genome sequences (mouse GRCm38/mm10) are downloaded from the GENCODE website (**Table S2**). Transcriptome sequences (cDNA sequences) and gene annotation downloaded from the same GENCODE source are used to search for putative ORFs. tRNA sequences are downloaded from UCSC Table Browser (53). rRNA sequences are downloaded from GENCODE (version M20, we have also tested M13 and M15) as well as published studies (18,19). The transcriptome is defined as the collection of all transcripts on the reference chromosomes. GENCODE Transcript biotypes are defined here – https://www.gencodegenes.org/pages/biotypes.html. In our pipeline, we remove the following biotypes:

- IG_* and TR_* (Immunoglobulin variable chain and T-cell receptor genes)
- miRNA
- misc_RNA
- Mt_rRNA and Mt_tRNA
- rRNA and ribozyme
- scaRNA, scRNA, snoRNA, snRNA and sRNA
- nonsense_mediated_decay
- Non_stop_decay

### Overview of strategy to identify candidate smORFs

Using the nucleotide sequences of all transcripts downloaded from GENCODE (release M13) (54) as a reference, ORFLine searches for putative ORFs beginning with “AUG”, “TUG”, “CUG”, “GUG” and ending with “UAG”, “UAA”, “UGA” in each of the three reading frames. It then removes ORFs that are not n*3 (n > 1) nucleotides long and designates those that are between 10 to 100 codons in length as putative smORFs. The ORF coordinates are initially transcript coordinates and are converted to genomic coordinates given exon location information in the gene annotation (in GTF/GFF format), the output of this step are genomic coordinates and strands for putative smORFs in BED format. Each ORF will be assigned two different identifiers, one is called RegionId, the second is called ORFId. RegionId is created based on genomic coordinates, ORFId is created based on the transcript coordinates. An ORF has a unique genomic location, thus RegionId is unique, but it may arise from multiple overlapping transcripts, so it may have multiple ORFIds (**Figure S1**). This step is carried out only once and needs to be updated when transcriptome annotation is changed.

### Sequencing data processing

Raw Illumina sequencing data in FASTQ format are trimmed of adapter sequences and the resultant reads are aligned against specific

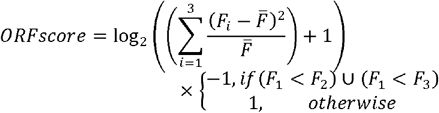

sequences assembled from a collection of rRNA, tRNA, Mt_rRNA and Mt_tRNA snRNA, snoRNA, misc_RNA and miRNA sequences using Bowtie v1.2.2 (55) to remove these sequences. The remaining reads are aligned to the reference genome (GRCm38). Adaptor trimming and quality trimming (including poor quality “N” base at the 5’ end of some of the reads) were performed with Trim Galore v0.4.5 (https://www.bioinformatics.babraham.ac.uk/projects/trim_galore), quality checked with FastQC v 0.11.8 (http://www.bioinformatics.bbsrc.ac.uk/projects/fastqc). Authentic RPFs will be ~28-30nt in length, therefore, we have kept trimmed reads that have a length between 25 and 35 nt, as they account for ~75% of the total reads on average (refer to information in **Table S1**).

#### Alignment to reference genome

The reads were mapped to the GRCm38/mm10 reference genome using the STAR aligner v2.5.2a (56). The aligner reports only uniquely mapped reads (mapping quality MAPQ = 255). The following shows an example command, and parameters are in bold:

STAR **--runThreadN** $THREAD \

**--genomeDir** $REFGENOMESTAR \
**--readFilesIn**$OUTPATH/bowtie-contanminant-removal/${NAME}_trimmed_unfiltered.fq.gz
**--readFilesCommand** zcat \
**--outReadsUnmapped** Fastx \
**--outFileNamePrefix** $OUTPATH/star-genome/$NAME/ \
**--alignIntronMin** $ALIGNINTRON_MIN \
**--alignIntronMax** $ALIGNINTRON_MAX \
**--alignEndsType** EndToEnd \
**--outFilterMismatchNmax** $MISMATCH_MAX \
**--outFilterMismatchNoverLmax** $MISMATCH_NOVERL_MAX \
**--outFilterType** $FILTER_TYPE \
**--outFilterIntronMotifs** RemoveNoncanonicalUnannotated \
**--outSAMattributes** $SAM_ATTR \
**--outSAMtype** BAM SortedByCoordinate \
**--outBAMsortingThreadN** $THREAD

#### Transcript expression quantification

In each experiment, sequence alignments (in BAM format) of all biological replicates were combined for RNA-Seq and Ribo-Seq respectively. Transcript expression was quantified using StringTie v1.3.6 (57) in FPKM (Fragments Per Kilobase per Million). From a given dataset, a minimal expression level was set to FPKM > 0.5 (27) to exclude nonexpressed transcripts.

#### P-site offset determination

A majority of RPFs has a length between 28-31 nucleotides (nt). The 5’ P-site offset is the distance from the 5’ end of the read to the ribosomal P-site (15). To determine P-site offset, we separated footprints into groups based on their lengths, P-site offset was estimated for each read length using plastid python library v0.4.8 (58). We observed P-site offsets are 12 nt long for RPF in 28-31 nt in our experiments.

### Identification of translated smORFs

ORFLine takes the gene annotation, putative smORFs, Ribo-Seq and RNA-Seq alignment as input to predict actively translated smORFs. ORFLine combines alignment files of all biological replicates (pooled analysis) to increase the signal intensity in case the smORFs are lowly expressed. This component consists of several metrics and filters, putative smORFs that have exceeded a confidence threshold for each metric (as indicated in **Table 1**) were kept.

#### RPF coverage

To filter ORFs which are insufficiently covered by reads, we calculated the proportion of codons being covered by RPFs. We consider a codon covered if there is a mapped RPF with the P-site aligned to nucleotide 1 of that codon. An ORF is discarded if the ratio of covered codons to the total number of codons in the ORF < 0.1 (18).

#### ORFScore

ORFScore was proposed by Bazzini and colleagues (18) and we reimplemented the ORFScore algorithm in R. The ORFScore was then calculated as:

where *F*_n_ is the number of reads in reading frame n, 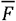 is the total number of reads across all three frames divided by 3. RPFs were counted at each position within an ORF, excluding the first and last coding codons. To filter out putative artifactual peaks, the most abundant read position was masked if reads aligning to that position comprised more than 70% of the total reads in the ORF. The ORFScore is a log-scaled chi-squared goodness of fit test statistic, the p-values associated with the test were adjusted using Benjamini-Hochberg FDR-controlling method and smORFs with ORFScore > 0 and adjusted p-value < 0.01 were retained.

#### Ribosome Release Score (RRS)

Firstly, the 3’UTRs of smORFs is defined. For canonical smORFs, we used annotated 3’UTRs. For other classes of smORFs, their 3’UTRs were defined as the region between the stop codon and the next possible start codon in any frame. The RRS score is defined as the ratio of the two normalized ratios and calculated with the following equation: RRS = (FPKM_RF ORF/FPKM_RF 3’-UTR)/ (FPKM_RNA ORF/FPKM_RNA 3’- UTR). Based on the original study, smORF with RRS > 5 is considered to be translated (26).

#### Inside/outside read ratio

The ribosome footprints typically show precise positioning between the start and the stop codon of translated ORFs. Low density of footprints before start codons and after stop codons and high inside/outside ratio is expected. By considering the read distribution of the nearest 3 upstream codons outside and the first 3 codons inside an ORF, we used a feature called inside/outside read ratio (total RPF of inside 3 codons/total RPF of outside 3 codons) to assess whether genuine translation takes place. ORFs will be discarded if the ratio ≤ 1 (more reads mapping outside than inside).

### Analysis of predicted smORFs

#### Translation efficiency (TE)

A measure of the rate of translation for a given feature (e.g. the CDS of a mRNA or a smORF), obtained in ribosome profiling experiments. It was calculated as the base 2 logarithmic ratio of RPF expression (FPKM) over mRNA expression (FPKM).

#### Conservation of the amino acid sequences

To examine the conservation of smORF-encoded micropeptide sequences between species, we performed PhyloCSF (59), a likelihood-based method to analyse signatures of evolutionary conservation in multiple species sequence alignments. PhyloCSF assigned a score to each smORF based on conservation within selected vertebrate species (https://github.com/mlin/PhyloCSF/wiki#available-phylogenies). For each smORF, we selected alignments of mouse, human, chimpanzee, gorilla, cow, dog and zebrafish from a publicly available whole genome multiple alignment using Galaxy “stitch gene blocks” tool (60). smORFs were considered conserved if their PhyloCSF score was > 50 (26), and weakly conserved if they had a PhyloCSF score > 0. PhyloCSF score = 0 indicates that there is no DNA sequence alignment cross species and PhyloCSF score < 0 is considered not conserved.

#### Gene ontology (GO) enrichment analysis

We used the g:Profiler server (61) to perform GO analysis in two unranked lists of genes mode. The background list comprised the combined expressed transcripts (FPKM > 0.5) of B and T cells. The target list contains the host transcripts of the smORFs. For the significance threshold, we chose the default option g:SCS threshold and the default value 0.05.

#### Secreted micropeptide prediction

We used the SignalP 4.0 server (40) to predict signal peptides present at the N-terminus of the micropeptide amino acid sequences. We used default parameters. For selected candidates, we ran prediction of transmembrane helices using the TMHMM 2.0 Server (62) (default parameters) to rule out transmembrane peptides.

#### Cloning and expression of candidate secreted micropeptides

The coding sequences of predicted smORFs were codon optimised for mammalian expression (GeneArt from ThermoFisher) and cloned into a mammalian expression vector. A C-terminal FLAG epitope tag was placed downstream of the micropeptide cDNA sequences flanked by a spacer sequence of Glycine-Alanine-Alanine. This is followed by an EMCV-IRES upstream of GFP cDNA. The di-cistronic mRNA is controlled by a CAG promoter. For transfection, HEK293T were seeded at 50% confluency the day before, and transfected at 60-90% confluency. TransIT (Mirus Bio) was used to deliver 1 μg plasmid per well of 6-well plates according to the manufacturer’s instruction. 3 hours post transfection, cells were replaced with fresh media to remove possible plasmid contamination in downstream assays. 21-24 hours post transfection, cells were collected by centrifugation at 300 × g for 5 minutes at 4 °C and lysed using RIPA buffer. Total secreted proteins were obtained by centrifugation of the culture media at 300 × g for 5 minutes at 4 °C. For Western blot analysis, total cell lysates and total supernatant were resolved with 16% Tris-Tricine/SDS-PAGE (63). After transfer, PVDF membranes were blotted with anti-FLAG (clone M2, Sigma) and anti-GFP (clone D5.1, Cell Signaling Technology) antibodies. Images were acquired and analysed using the Odyssey CLx (LI-COR).

## End Matter

### Data Availability

| Cell type | RNA-Seq | Ribo-Seq |
| --- | --- | --- |
| B cell setup 1 | GSE62129 | GSE62134 |
| B cell setup 2 | GSE146073<br>(GSM4364117-121) | GSE154491 (all samples) |
| CD4+ T cell | GSE155087<br>(GSM4694962-64) | GSE155087<br>(GSM4694968-972) |
| Th1 reactivation | GSE83351 | GSE83351 |

### Pipeline Availability

Pipeline code is publicly available on the source code hosting platform GitHub. The URL is https://github.com/boboppie/ORFLine. We also created a Singularity image (https://singularity.lbl.gov/) which enables the users to execute and test the pipeline easily in a virtual environment. All dependencies including bioinformatics tools are pre-installed in the image, the URL is https://github.com/boboppie/ORFLine-singularity.

### Supplementary Data

Supplementary Data are available at BioRxiv online.

## Acknowledgments

We thank Sebastian Lukasiak for sharing datasets for analysis. We thank Louise Matheson, Joanna Krupka and Daniel Hodson for critical reading of the manuscript. We thank Simon Andrews and Joanna Krupka for testing ORFLine and sharing their valuable feedback to improve the code. Public RNA-Seq datasets (**Table S6**) used in this study were processed by Babraham Bioinformatics group. We thank Finkelstein Lab (https://github.com/finkelsteinlab) for the BioRxiv template.

## References

1. Makarewich, C.A. and Olson, E.N. (2017) Mining for Micropeptides. Trends in cell biology, 27, 685–696.

2. Yin, X., Jing, Y. and Xu, H. (2019) Mining for missed sORF-encoded peptides. Expert Rev Proteomics, 16, 257–266.

3. Orr, M.W., Mao, Y., Storz, G. and Qian, S.B. (2020) Alternative ORFs and small ORFs: shedding light on the dark proteome. Nucleic Acids Res, 48, 1029–1042.

4. Couso, J.P. and Patraquim, P. (2017) Classification and function of small open reading frames. Nat Rev Mol Cell Biol, 18, 575–589.

5. Kondo, T., Hashimoto, Y., Kato, K., Inagaki, S., Hayashi, S. and Kageyama, Y. (2007) Small peptide regulators of actin-based cell morphogenesis encoded by a polycistronic mRNA. Nat Cell Biol, 9, 660–665.

6. Slavoff, S.A., Heo, J., Budnik, B.A., Hanakahi, L.A. and Saghatelian, A. (2014) A human short open reading frame (sORF)-encoded polypeptide that stimulates DNA end joining. J Biol Chem, 289, 10950–10957.

7. D’Lima, N.G., Ma, J., Winkler, L., Chu, Q., Loh, K.H., Corpuz, E.O., Budnik, B.A., Lykke-Andersen, J., Saghatelian, A. and Slavoff, S.A. (2017) A human microprotein that interacts with the mRNA decapping complex. Nat Chem Biol, 13, 174–180.

8. Magny, E.G., Pueyo, J.I., Pearl, F.M., Cespedes, M.A., Niven, J.E., Bishop, S.A. and Couso, J.P. (2013) Conserved regulation of cardiac calcium uptake by peptides encoded in small open reading frames. Science, 341, 1116–1120.

9. Lee, C., Zeng, J., Drew, B.G., Sallam, T., Martin-Montalvo, A., Wan, J., Kim, S.J., Mehta, H., Hevener, A.L., de Cabo, R. et al. (2015) The mitochondrial-derived peptide MOTS-c promotes metabolic homeostasis and reduces obesity and insulin resistance. Cell Metab, 21, 443–454.

10. Matsumoto, A., Pasut, A., Matsumoto, M., Yamashita, R., Fung, J., Monteleone, E., Saghatelian, A., Nakayama, K.I., Clohessy, J.G. and Pandolfi, P.P. (2017) mTORC1 and muscle regeneration are regulated by the LINC00961-encoded SPAR polypeptide. Nature, 541, 228–232.

11. Huang, J.Z., Chen, M., Chen, Gao, X.C., Zhu, S., Huang, H., Hu, M., Zhu, H. and Yan, G.R. (2017) A Peptide Encoded by a Putative lncRNA HOXB-AS3 Suppresses Colon Cancer Growth. Mol Cell, 68, 171–184 e176.

12. Jackson, R., Kroehling, L., Khitun, A., Bailis, W., Jarret, A., York, A.G., Khan, O.M., Brewer, J.R., Skadow, M.H., Duizer, C. et al. (2018) The translation of non-canonical open reading frames controls mucosal immunity. Nature, 564, 434–438.

13. Carninci, P., Kasukawa, T., Katayama, S., Gough, J., Frith, M. C., Maeda, N., Oyama, R., Ravasi, T., Lenhard, B., Wells, C. et al. (2005) The transcriptional landscape of the mammalian genome. Science, 309, 1559–1563.

14. Ingolia, N.T., Brar, G.A., Stern-Ginossar, N., Harris, M.S., Talhouarne, G.J., Jackson, S.E., Wills, M.R. and Weissman, J.S. (2014) Ribosome profiling reveals pervasive translation outside of annotated protein-coding genes. Cell reports, 8, 1365–1379.

15. Ingolia, N.T., Ghaemmaghami, S., Newman, J.R. and Weissman, J.S. (2009) Genome-wide analysis in vivo of translation with nucleotide resolution using ribosome profiling. Science, 324, 218–223.

16. Ingolia, N.T., Lareau, L.F. and Weissman, J.S. (2011) Ribosome profiling of mouse embryonic stem cells reveals the complexity and dynamics of mammalian proteomes. Cell, 147, 789–802.

17. Dunn, J.G., Foo, C.K., Belletier, N.G., Gavis, E.R. and Weissman, J.S. (2013) Ribosome profiling reveals pervasive and regulated stop codon readthrough in Drosophila melanogaster. Elife, 2, e01179.

18. Bazzini, A.A., Johnstone, T.G., Christiano, R., Mackowiak, S.D., Obermayer, B., Fleming, E.S., Vejnar, C.E., Lee, M.T., Rajewsky, N., Walther, T.C. et al. (2014) Identification of small ORFs in vertebrates using ribosome footprinting and evolutionary conservation. The EMBO journal, 33, 981–993.

19. Fields, A.P., Rodriguez, E.H., Jovanovic, M., Stern-Ginossar, N., Haas, B.J., Mertins, P., Raychowdhury, R., Hacohen, N., Carr, S.A., Ingolia, N.T. et al. (2015) A Regression-Based Analysis of Ribosome-Profiling Data Reveals a Conserved Complexity to Mammalian Translation. Mol Cell, 60, 816–827.

20. Ji, Z., Song, R., Regev, A. and Struhl, K. (2015) Many lncRNAs, 5’UTRs, and pseudogenes are translated and some are likely to express functional proteins. eLife, 4, e08890.

21. Johnstone, T.G., Bazzini, A.A. and Giraldez, A.J. (2016) Upstream ORFs are prevalent translational repressors in vertebrates. The EMBO journal, 35, 706–723.

22. Calviello, L., Mukherjee, N., Wyler, E., Zauber, H., Hirsekorn, A., Selbach, M., Landthaler, M., Obermayer, B. and Ohler, U. (2016) Detecting actively translated open reading frames in ribosome profiling data. Nat Methods, 13, 165–170.

23. Malone, B., Atanassov, I., Aeschimann, F., Li, X., Grosshans, H. and Dieterich, C. (2017) Bayesian prediction of RNA translation from ribosome profiling. Nucleic Acids Res, 45, 2960–2972.

24. Xiao, Z., Huang, R., Xing, X., Chen, Y., Deng, H. and Yang, X. (2018) De novo annotation and characterization of the translatome with ribosome profiling data. Nucleic Acids Res, 46, e61.

25. Liu, Q., Shvarts, T., Sliz, P. and Gregory, R.I. (2020) RiboToolkit: an integrated platform for analysis and annotation of ribosome profiling data to decode mRNA translation at codon resolution. Nucleic Acids Res.

26. Guttman, M., Russell, P., Ingolia, N.T., Weissman, J.S. and Lander, E.S. (2013) Ribosome profiling provides evidence that large noncoding RNAs do not encode proteins. Cell, 154, 240–251.

27. Hart, T., Komori, H.K., LaMere, S., Podshivalova, K. and Salomon, D.R. (2013) Finding the active genes in deep RNA-seq gene expression studies. BMC Genomics, 14, 778.

28. Diaz-Munoz, M.D., Bell, S.E., Fairfax, K., Monzon-Casanova, E., Cunningham, A.F., Gonzalez-Porta, M., Andrews, S.R., Bunik, V.I., Zarnack, K., Curk, T. et al. (2015) The RNA-binding protein HuR is essential for the B cell antibody response. Nat Immunol, 16, 415–425.

29. Davari, K., Lichti, J., Gallus, C., Greulich, F., Uhlenhaut, N.H., Heinig, M., Friedel, C.C. and Glasmacher, E. (2017) Rapid Genome-wide Recruitment of RNA Polymerase II Drives Transcription, Splicing, and Translation Events during T Cell Responses. Cell reports, 19, 643–654.

30. Michel, A.M., Choudhury, K.R., Firth, A.E., Ingolia, N.T., Atkins, J.F. and Baranov, P.V. (2012) Observation of dually decoded regions of the human genome using ribosome profiling data. Genome Res, 22, 2219–2229.

31. Na, C.H., Barbhuiya, M.A., Kim, M.S., Verbruggen, S., Eacker, S.M., Pletnikova, O., Troncoso, J.C., Halushka, M.K., Menschaert, G., Overall, C.M. et al. (2018) Discovery of noncanonical translation initiation sites through mass spectrometric analysis of protein N termini. Genome Res, 28, 25–36.

32. Sha, J., Zhao, G., Chen, X., Guan, W., He, Y. and Wang, Z. (2012) Antibacterial potential of hGlyrichin encoded by a human gene. J Pept Sci, 18, 97–104.

33. Andrews, S.J. and Rothnagel, J.A. (2014) Emerging evidence for functional peptides encoded by short open reading frames. Nat Rev Genet, 15, 193–204.

34. Calvo, S.E., Pagliarini, D.J. and Mootha, V.K. (2009) Upstream open reading frames cause widespread reduction of protein expression and are polymorphic among humans. Proc Natl Acad Sci U S A, 106, 7507–7512.

35. Wang, X.Q. and Rothnagel, J.A. (2004) 5’-untranslated regions with multiple upstream AUG codons can support low-level translation via leaky scanning and reinitiation. Nucleic Acids Res, 32, 1382–1391.

36. Chew, G.L., Pauli, A. and Schier, A.F. (2016) Conservation of uORF repressiveness and sequence features in mouse, human and zebrafish. Nature communications, 7, 11663.

37. Zhang, H., Wang, Y. and Lu, J. (2019) Function and Evolution of Upstream ORFs in Eukaryotes. Trends Biochem Sci, 44, 782–794.

38. Altschul, S.F., Gish, W., Miller, W., Myers, E.W. and Lipman, D.J. (1990) Basic local alignment search tool. J Mol Biol, 215, 403–410.

39. Gunisova, S., Hronova, V., Mohammad, M.P., Hinnebusch, A.G. and Valasek, L.S. (2018) Please do not recycle! Translation reinitiation in microbes and higher eukaryotes. FEMS Microbiol Rev, 42, 165–192.

40. Petersen, T.N., Brunak, S., von Heijne, G. and Nielsen, H. (2011) SignalP 4.0: discriminating signal peptides from transmembrane regions. Nat Methods, 8, 785–786.

41. Bhatta, A., Atianand, M., Jiang, Z., Crabtree, J., Blin, J. and Fitzgerald, K.A. (2020) A Mitochondrial Micropeptide Is Required for Activation of the Nlrp3 Inflammasome. J Immunol, 204, 428–437.

42. Shi, W., Liao, Y., Willis, S.N., Taubenheim, N., Inouye, M., Tarlinton, D.M., Smyth, G.K., Hodgkin, P.D., Nutt, S.L. and Corcoran, L.M. (2015) Transcriptional profiling of mouse B cell terminal differentiation defines a signature for antibody-secreting plasma cells. Nat Immunol, 16, 663–673.

43. Luo, C.T., Liao, W., Dadi, S., Toure, A. and Li, M.O. (2016) Graded Foxo1 activity in Treg cells differentiates tumour immunity from spontaneous autoimmunity. Nature, 529, 532–536.

44. Sendoel, A., Dunn, J.G., Rodriguez, E.H., Naik, S., Gomez, N.C., Hurwitz, B., Levorse, J., Dill, B.D., Schramek, D., Molina, H. et al. (2017) Translation from unconventional 5’ start sites drives tumour initiation. Nature, 541, 494–499.

45. Zhang, S., Reljic, B., Liang, C., Kerouanton, B., Francisco, J.C., Peh, J.H., Mary, C., Jagannathan, N.S., Olexiouk, V., Tang, C. et al. (2020) Mitochondrial peptide BRAWNIN is essential for vertebrate respiratory complex III assembly. Nature communications, 11, 1312.

46. Murat, P., Marsico, G., Herdy, B., Ghanbarian, A., Portella, G. and Balasubramanian, S. (2018) RNA G-quadruplexes at upstream open reading frames cause DHX36- and DHX9- dependent translation of human mRNAs. Genome Biology, 19, 229.

47. Wu, Q., Wright, M., Gogol, M.M., Bradford, W.D., Zhang, N. and Bazzini, A.A. (2020) Translation of small downstream ORFs enhances translation of canonical main open reading frames. The EMBO journal, e104763.

48. Oyama, M., Kozuka-Hata, H., Suzuki, Y., Semba, K., Yamamoto, T. and Sugano, S. (2007) Diversity of translation start sites may define increased complexity of the human short ORFeome. Mol Cell Proteomics, 6, 1000–1006.

49. Slavoff, S.A., Mitchell, A.J., Schwaid, A.G., Cabili, M.N., Ma, J., Levin, J.Z., Karger, A.D., Budnik, B.A., Rinn, J.L. and Saghatelian, A. (2013) Peptidomic discovery of short open reading frame–encoded peptides in human cells. Nature Chemical Biology, 9, 59–64.

50. Chen, J., Brunner, A.D., Cogan, J.Z., Nunez, J.K., Fields, A.P., Adamson, B., Itzhak, D.N., Li, J.Y., Mann, M., Leonetti, M.D. et al. (2020) Pervasive functional translation of noncanonical human open reading frames. Science, 367, 1140–1146.

51. Dersh, D., Holly, J. and Yewdell, J.W. (2020) A few good peptides: MHC class I-based cancer immunosurveillance and immunoevasion. Nat Rev Immunol.

52. Harrow, J., Frankish, A., Gonzalez, J.M., Tapanari, E., Diekhans, M., Kokocinski, F., Aken, B.L., Barrell, D., Zadissa, A., Searle, S. et al. (2012) GENCODE: the reference human genome annotation for The ENCODE Project. Genome Res, 22, 1760–1774.

53. Karolchik, D., Hinrichs, A.S., Furey, T.S., Roskin, K.M., Sugnet, C.W., Haussler, D. and Kent, W.J. (2004) The UCSC Table Browser data retrieval tool. Nucleic Acids Res, 32, D493–496.

54. Frankish, A., Uszczynska, B., Ritchie, G.R., Gonzalez, J.M., Pervouchine, D., Petryszak, R., Mudge, J.M., Fonseca, N., Brazma, A., Guigo, R. et al. (2015) Comparison of GENCODE and RefSeq gene annotation and the impact of reference geneset on variant effect prediction. BMC Genomics, 16 Suppl 8, S2.

55. Langmead, B., Trapnell, C., Pop, M. and Salzberg, S.L. (2009) Ultrafast and memory-efficient alignment of short DNA sequences to the human genome. Genome Biol, 10, R25.

56. Dobin, A., Davis, C.A., Schlesinger, F., Drenkow, J., Zaleski, C., Jha, S., Batut, P., Chaisson, M. and Gingeras, T.R. (2013) STAR: ultrafast universal RNA-seq aligner. Bioinformatics, 29, 15–21.

57. Pertea, M., Pertea, G.M., Antonescu, C.M., Chang, T.C., Mendell, J.T. and Salzberg, S.L. (2015) StringTie enables improved reconstruction of a transcriptome from RNA-seq reads. Nature biotechnology, 33, 290–295.

58. Dunn, J.G. and Weissman, J.S. (2016) Plastid: nucleotide-resolution analysis of next-generation sequencing and genomics data. BMC Genomics, 17, 958.

59. Lin, M.F., Jungreis, I. and Kellis, M. (2011) PhyloCSF: a comparative genomics method to distinguish protein coding and non-coding regions. Bioinformatics, 27, i275–282.

60. Blankenberg, D., Taylor, J., Nekrutenko, A. and Galaxy, T. (2011) Making whole genome multiple alignments usable for biologists. Bioinformatics, 27, 2426–2428.

61. Raudvere, U., Kolberg, L., Kuzmin, I., Arak, T., Adler, P., Peterson, H. and Vilo, J. (2019) g:Profiler: a web server for functional enrichment analysis and conversions of gene lists (2019 update). Nucleic Acids Res, 47, W191–W198.

62. Krogh, A., Larsson, B., von Heijne, G. and Sonnhammer, E.L. (2001) Predicting transmembrane protein topology with a hidden Markov model: application to complete genomes. J Mol Biol, 305, 567–580.

63. Haider, S.R., Reid, H.J. and Sharp, B.L. (2019) Tricine-SDS-PAGE. Methods in molecular biology, 1855, 151–160.

