## Supplementary Data for "ORFLine: a bioinformatic pipeline to prioritise small open reading frames identifies candidate secreted small proteins from lymphocytes"

**Supplemental tables and figures:**

**Table S1.** Sequencing metrics.

**Table S2.** Reference annotation. List of public sequences and annotation files used in the pipeline. Reference genome and transcriptome sequences, gene annotation (mouse and human) are from GENCODE. Ribosomal RNA (rRNA) and Transfer RNA (tRNA) sequences are from UCSC Table Browser. rRNA sequences are from Ensembl.

**Table S3.** Predicted smORFs. The columns are Region ID, Chromosome, Start, Stop, Strand, Class, Transcript ID (Ensembl), Gene symbol, Gene description, PhyloCSF score, AA length, Peptide sequence, Cell types.

**Table S4.** GO enrichment.

**Table S5.** dORF conservation.

**Table S6.** Public datasets used in this study.

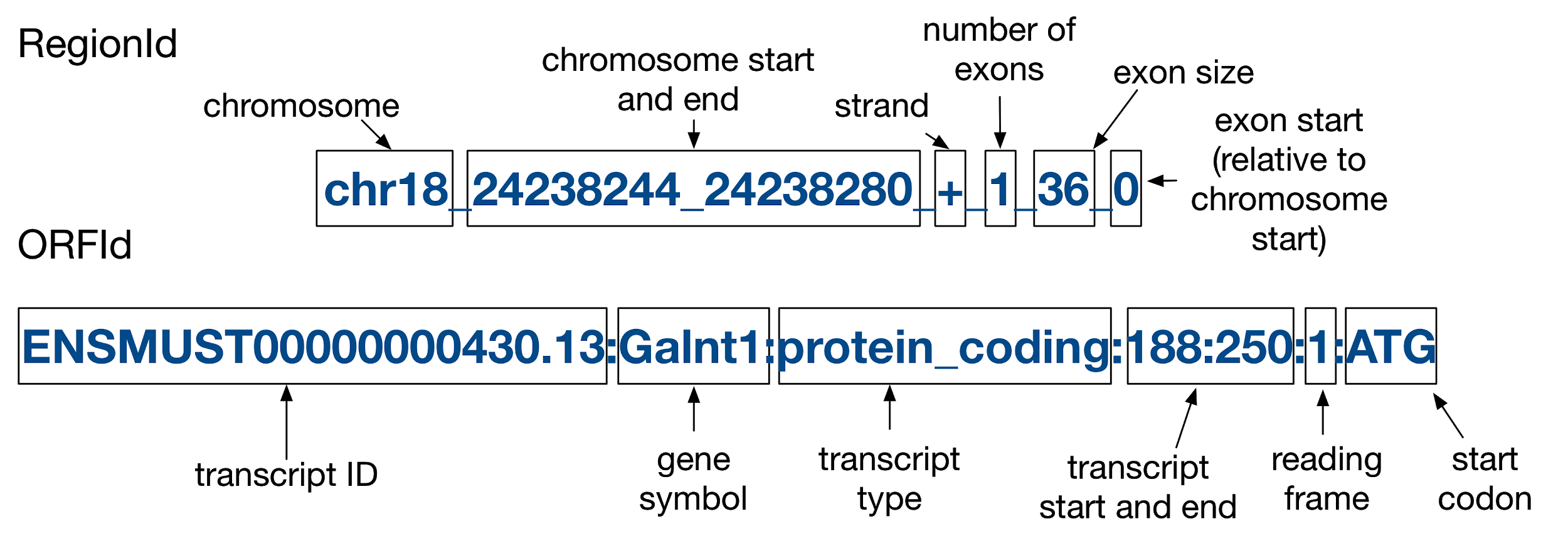

**Figure S1.** RegionId and ORFId explained. RegionId is genomic-based, it indicates the unique location of a smORF on the genome. ORFId is transcript-based, it contains information regarding the smORF’s relative position on its host transcript.

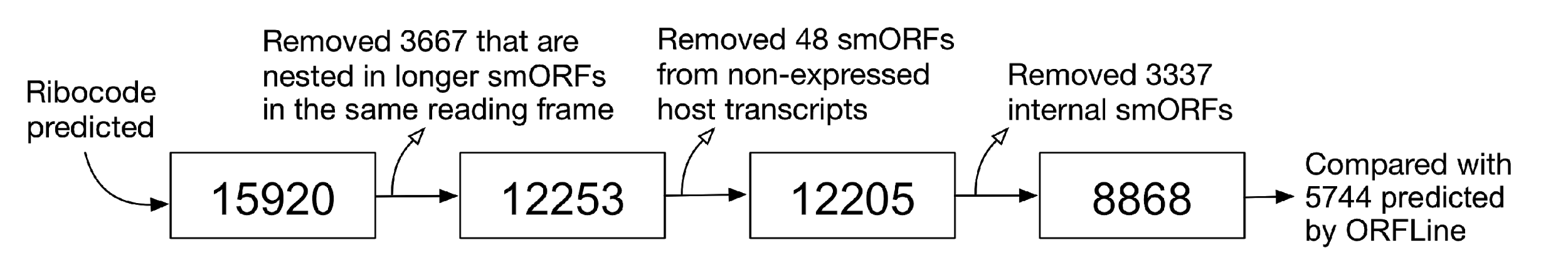

**Figure S2**. Number of smORFs used for a comparison between RiboCode and ORFLine. Initially, 15920 smORFs were predicted by RiboCode, 3367 were removed as they were nested in longer smORFs in the same frame. 48 were removed as they were from non-expressed host transcripts, and 3337 were removed as they were internal smORFs. The remaining 8868 were used to compare with ORFLine prediction.

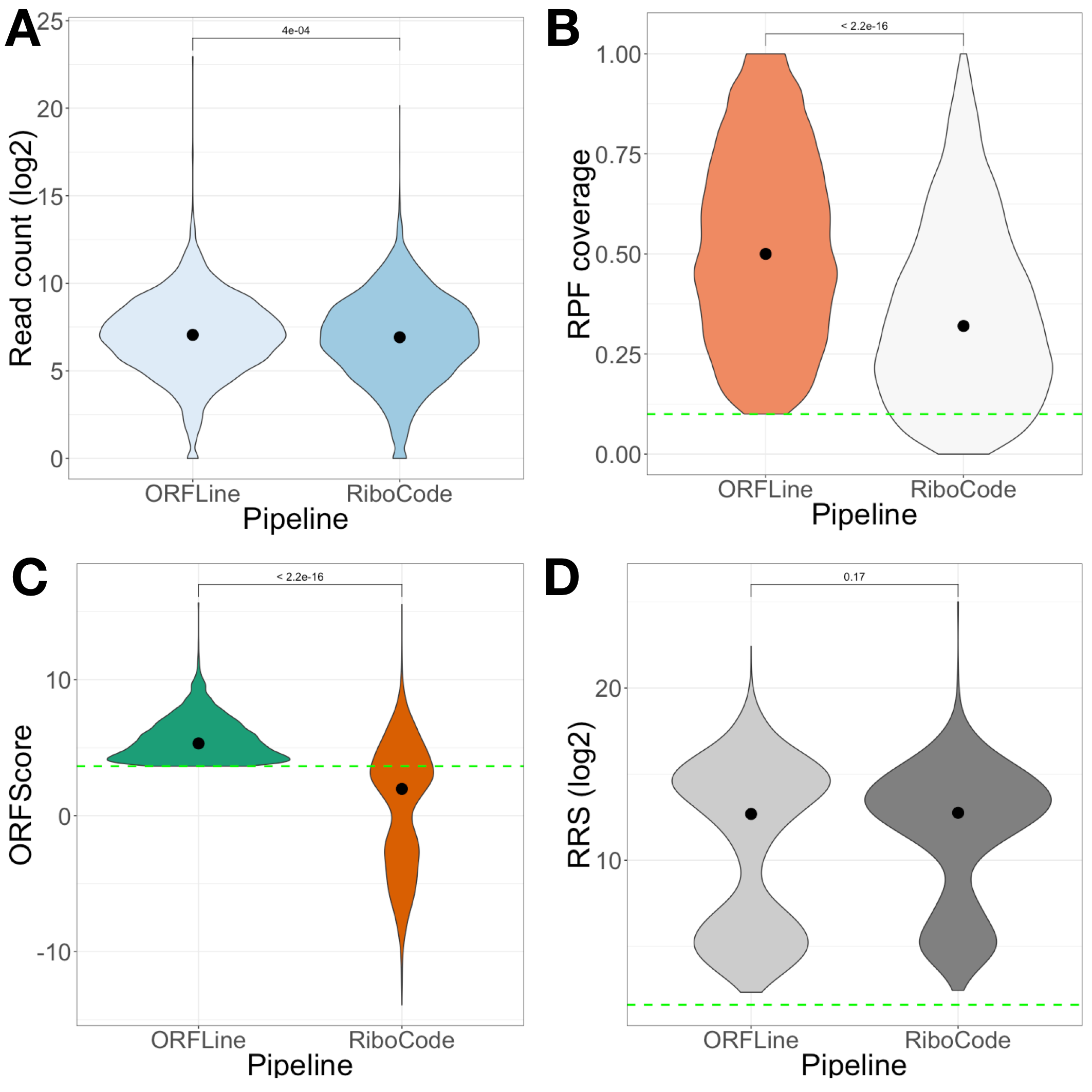

**Figure S3.** ORFLine predicts putative smORFs with more robust metrics. The following metrics of smORFs differentially predicted by ORFLine and RiboCode were compared: A) Read count B) RPF coverage C) ORFScore and D) RRS. Green dotted lines showed the threshold used by ORFLine in the according metrics.

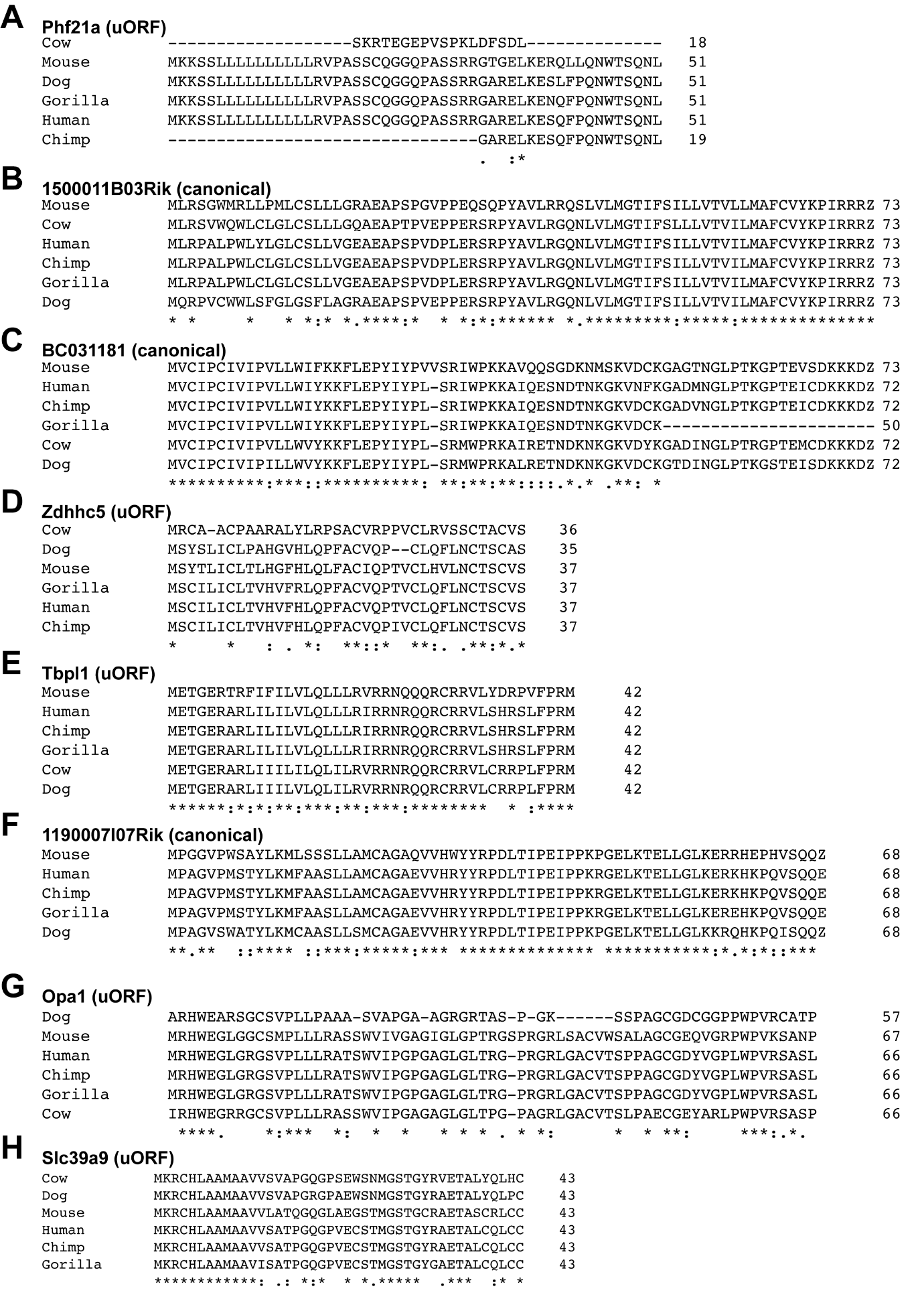

**Figure S4.** Selected putative micropeptides are conserved by amino acid sequence. The amino acid sequence of the following micropeptides from mouse, human, chimpanzee, gorilla, cow and dog are compared: A) Phf21a (uORF) B) 1500011B03Rik C) BC031181 D) Zdhhc5 (uORF) E) Tbpl1 (uORF) F) 1190007I07Rik G) Opa1 (uORF) H) Slc39a9 (uORF). Conservation was displayed as followed: * fully conserved : strongly similar . weakly similar.
